# O-Linked Sialoglycans Modulate the Proteolysis of SARS-CoV-2 Spike and Likely Contribute to the Mutational Trajectory in Variants of Concern

**DOI:** 10.1101/2022.09.15.508093

**Authors:** Edgar Gonzalez-Rodriguez, Mia Zol-Hanlon, Ganka Bineva-Todd, Andrea Marchesi, Mark Skehel, Keira E. Mahoney, Chloë Roustan, Annabel Borg, Lucia Di Vagno, Svend Kjaer, Antoni G. Wrobel, Donald J. Benton, Philipp Nawrath, Sabine L. Flitsch, Dhira Joshi, Andrés Manuel González-Ramírez, Katalin A. Wilkinson, Robert J. Wilkinson, Emma C. Wall, Ramón Hurtado-Guerrero, Stacy A. Malaker, Benjamin Schumann

## Abstract

The emergence of a polybasic cleavage motif for the protease furin in the SARS-CoV-2 spike protein has been established as a major factor for enhanced viral transmission in humans. The peptide region N-terminal to that motif is extensively mutated in major variants of concern including Alpha, Delta and Omicron. Besides furin, spike proteins from these variants appear to rely on other proteases for maturation, including TMPRSS2 that may share the same cleavage motif. Glycans found near the cleavage site have raised questions about proteolytic processing and the consequences of variant-borne mutations. Here, with a suite of chemical tools, we establish O-linked glycosylation as a major determinant of SARS-CoV-2 spike cleavage by the host proteases furin and TMPRSS2, and as a likely driving force for the emergence of common mutations in variants of concern. We provide direct evidence that the glycosyltransferase GalNAc-T1 primes glycosylation at Thr678 in the living cell, and this glycosylation event is suppressed by many, but not all variant mutations. A novel strategy for rapid bioorthogonal modification of Thr678-containing glycopeptides revealed that introduction of a negative charge completely abrogates furin activity. In a panel of synthetic glycopeptides containing elaborated O-glycans, we found that the sole incorporation of N-acetylgalactosamine did not substantially impact furin activity, but the presence of sialic acid in elaborated O-glycans reduced furin rate by up to 65%. Similarly, O-glycosylation with a sialylated trisaccharide had a negative impact on spike cleavage by TMPRSS2. With a chemistry-centered approach, we firmly establish O-glycosylation as a major determinant of spike maturation and propose that a disruption of O-GalNAc glycosylation is a substantial driving force for the evolution of variants of concern.

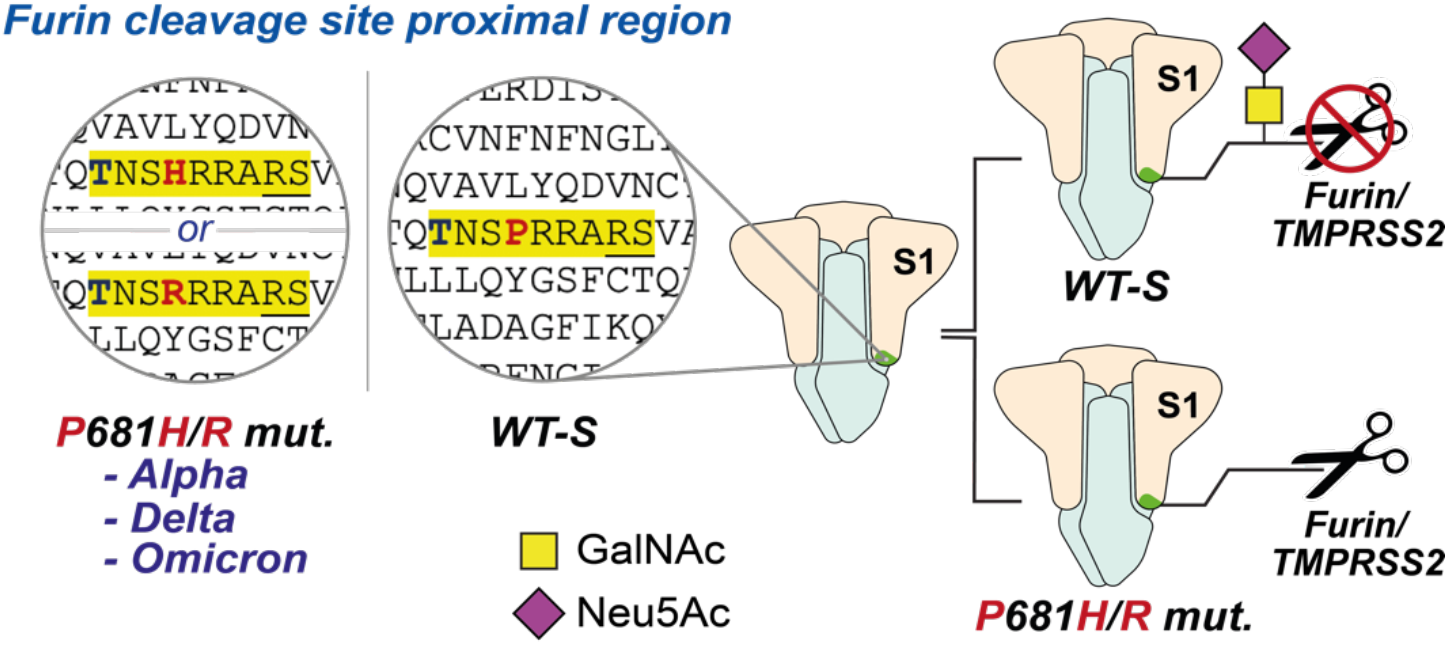

## MAIN

The viral surface spike protein has been the subject of intense scientific efforts to understand and curb SARS-CoV-2 transmission.^1–14^ Spike is a trimeric, multidomain glycoprotein (**Figure 1a**), with a dense glycan coat that plays crucial structural, immunological and functional roles.^15–28^ An evolutionarily novel arginine-rich peptide sequence in SARS-CoV-2 spike has been identified as the cleavage site for the Golgi-localised convertase furin. This furin cleavage site (FCS) is crucial for SARS-CoV-2 transmission, as cleavage enhances receptor binding and likely the fusion activity of spike.^3,29–31^ On a molecular level, furin hydrolyses the peptide bond between Arg685 and Ser686, converting full-length spike (termed FL-S) into the fragments S1 and S2 in the mature protein.^7,32–39^ Another host protease, TMPRSS2, has been proposed to act synergistically with furin, potentially also targeting the FCS with preference to cleave before arginine.^40,41^

**Figure 1.**
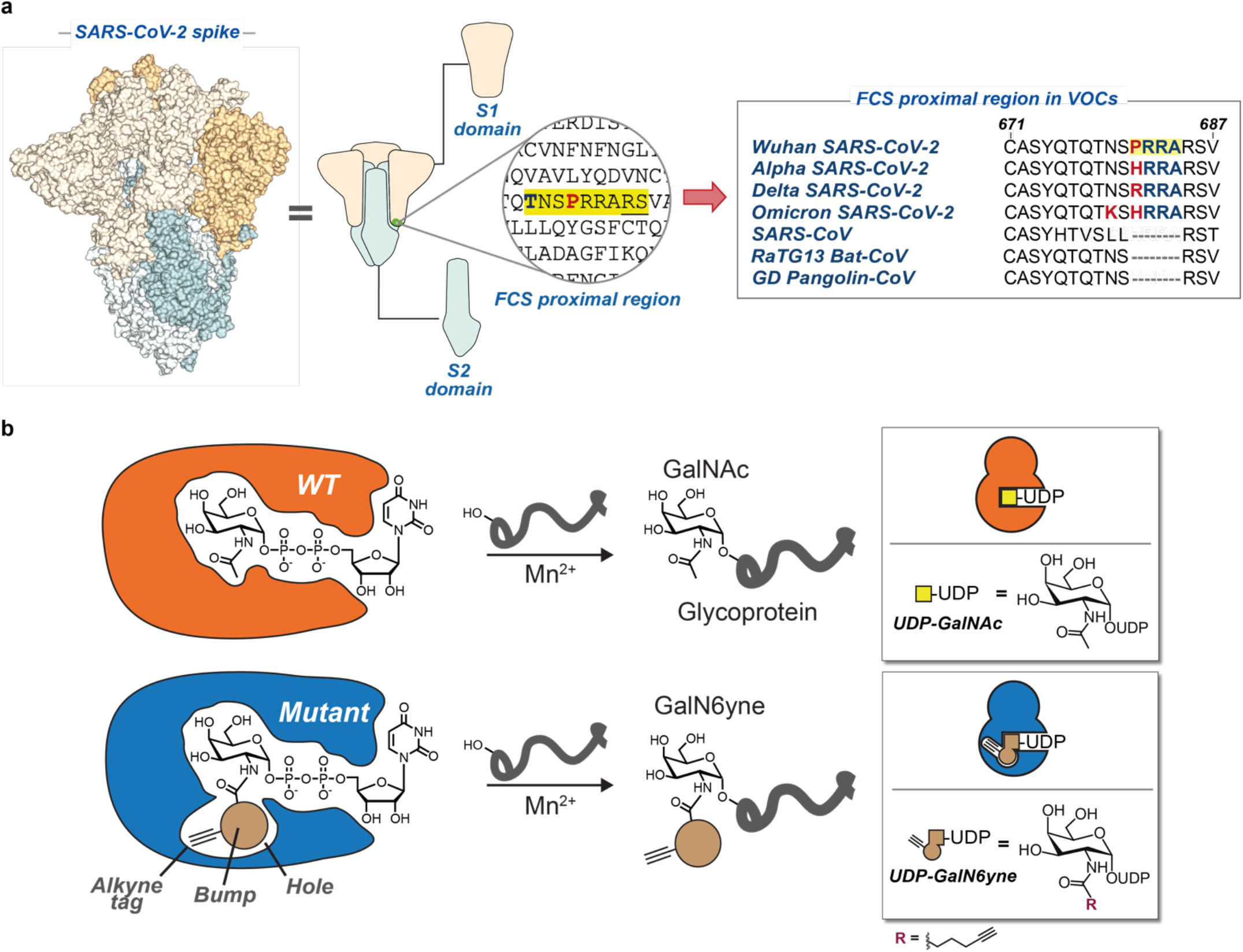
Dissecting O-glycosylation on SARS-CoV2 Spike. **a)** *Left*: SARS-CoV-2 spike model (6ZGE) and (*middle*) its corresponding cartoon representation with the furin cleavage site (FCS) proximal region highlighted in yellow. The blue and bold T corresponds to Thr678 which is a potential glycosylation site within the FCS proximal region. Underlined R and S residues correspond to the FCS. The S1 domain (including Thr678) lies N-terminal to the FCS and the S2 domain is C-terminal to the FCS. *Right*: peptide alignment for SARS-CoV-2 variants of concern (VOCs) and related coronaviruses showing the emergence of the polybasic motif in the FCS proximal region. Highlighted in yellow is the polybasic motif of SARS-CoV-2 spike. Bold and red are amino acid changes in positions 679 and 681 in VOCs. **b)** Bump-and-hole engineering allows for GalNAc-T isoenzyme-specific tagging of glycosylation substrates using the clickable substrate UDP-GalN6yne. FCS = furin cleavage site; VOCs = variants of concern.

Circulating variants of concern (VOCs) display increased proteolytic processing of spike into S1/S2.^37,42–44^ This increase has been associated with a remarkable polymorphism in the peptide stretch preceding the FCS between residues Gln675 and Pro681. Most VOCs and many minor circulating variants carry at least one mutation in that peptide region: Alpha (B.1.1.7) and Delta (B.1.617.2) display mutations at Pro681 to His and Arg, respectively, whereas all Omicron sub-lineages including BA.1, BA.2 and BA.5 combine P681H with the mutation N679K. Mutations in this region appear to have arisen more than once independently: Delta (P681R), Alpha (P681H) and Omicron lineages (P681H) are on different arms of the evolutionary tree with common ancestors that do not contain mutations around the FCS, indicating that some selection pressure on this sequence must have been present during their evolution.^45,46^ Lower-prominence variants have featured substitutions at Gln675 and Gln677,^47–51^ usually to amino acids with basic functionalities (**Figure 1a**). In line with the cleavage enhancing effect of these mutations, preparations of VOC spike from eukaryotic expression systems contain less FL-S than WT (Wuhan/WH04/2020) spike preparations.^52^ While it is tempting to suggest that an increase in basic amino acids enhances furin activity simply due to electrostatic extension of the polybasic cleavage site,^53,54^ this amino acid-centric notion neglects the impact of post-translational modifications. In line with this notion, Whittaker and colleagues observed that the P681H mutation alone does not increase furin cleavage of synthetic peptides.^55^ Due to the importance of spike in the viral infectious cycle, the key determinants of processing offer essential insights into the cell biology of viral maturation.

Like most surface proteins on animal viruses, SARS-CoV-2 spike is extensively coated with glycans that impact the mature virus’ infectivity and immunogenicity.^15–28^ Among these, Asn (N)-linked glycosylation is straightforward to predict due to the existence of a peptide consensus sequence (N-X-S/T; where X = any amino acid except Pro). In contrast, the prediction of Ser/Thr-linked N-acetylgalactosamine (O-GalNAc) glycosylation, which also greatly impacts viral biology,^25,56–60^ is an analytical challenge due to greater biosynthetic complexity and the lack of a peptide consensus sequence.^61–71^ Notably, the peptide region between Gln675 and Pro681 of SARS-CoV-2 spike harbours multiple Ser/Thr residues that may carry O-GalNAc glycans.^19,26,27^ Despite the analytical difficulties in understanding O-glycan biology, emerging data suggests that O-GalNAc glycosylation impacts furin-mediated spike cleavage.^72^ Due to the relevance of furin cleavage for viral infectivity, understanding the role of glycosylation in this process is essential.

The biosynthesis of O-GalNAc glycans is initiated by the introduction of the sugar N-acetylgalactosamine (GalNAc) from activated substrate uridine diphosphate (UDP)-GalNAc on to Ser/Thr side chains by a family of 20 GalNAc transferase (GalNAc-T1…T20) isoenzymes. GalNAc-Ts are often associated with isoenzyme-specific, decisive roles in physiological processes that are beginning to be unraveled.^73–82^ Understanding the substrate profiles of individual GalNAc-T isoenzymes yields insight into the regulation of such processes and can be the basis for the development of tools, diagnostics and therapeutics. However, assigning glycosylation sites to individual GalNAc-Ts is challenging due to their complex and often overlapping interplay in the secretory pathway.^83^ Additionally, the initial GalNAc residue is often further elaborated, generating mature glycans containing galactose (Gal), N-acetylglucosamine (GlcNAc), and the acidic monosaccharide N-acetylneuraminic acid (Neu5Ac) as a capping structure, further complicating the analytical profiling of O-GalNAc glycoproteins by mass spectrometry (MS) glycoproteomics. Indirect methods are thus often necessary to establish links between GalNAc-T isoenzymes and the glycosylation sites they modify to yield insights into O-glycan biology.^73,84–87^ Through co-expression of the individual human GalNAc-Ts with spike in insect cells and lectin staining, Ten Hagen and colleagues identified seven isoenzymes capable of introducing GalNAc into recombinant spike.^72^ Furthermore, co-expression of spike with GalNAc-T1 decreased furin cleavage, suggesting that glycosylation may modulate furin cleavage and, by implication, viral infectivity. This is in line with earlier findings that furin cleavage of other secreted proteins can be impacted by O-glycosylation.^88–92^ However, the biosynthetic complexity and technical challenges associated with O-glycoproteome analysis have thus far hindered closer investigation. Specifically, we currently lack knowledge on glycan abundance, precise attachment site(s), structural influence on proteolysis, and the roles of VOC mutations on O-GalNAc glycosylation.

By enabling a more direct view into the details of glycan biosynthesis, chemical tools have provided an insight into glycobiology that is orthogonal to classical methods of molecular biology. For example, by using a tactic termed “bump-and-hole engineering”, we have developed a chemical reporter strategy for the activities of individual GalNAc-T isoenzymes in the living cell (**Figure 1b**).^93,94^ Through structure-based design, the active site of a GalNAc-T isoenzyme was expanded by mutagenesis to contain a “hole”, which is complementary to a “bump” in a chemically modified analogue of the substrate UDP-GalNAc.^94^ The bumped substrate “UDP-GalN6yne” contained an alkyne tag that enabled the bioorthogonal ligation of fluorophores or biotin after transfer to a glycoprotein, allowing the profiling of the substrates of individual GalNAc-Ts.^94–96^ Recently, we introduced a clickable, positively charged imidazolium tag (termed ITag) that enhances MS-based analysis by increasing the charge state and improving the fragmentation-based sequencing of glycopeptides.^97^ Importantly, UDP-GalN6yne can be biosynthesized in the living cell through the introduction of an artificial metabolic pathway and feeding with a membrane-permeable peracetylated GalN6yne precursor (Ac_4_GalN6yne), allowing for the installation of a fully functional GalNAc-T bump-and-hole system.^94,98^ Building on the power of our chemical tools to dissect the role of O-GalNAc glycosylation, we sought to map the molecular details of glycan-mediated modulation of spike processing.

Here, with aid from this repertoire of chemical biology tools, we establish O-linked glycosylation as a major determinant of SARS-CoV-2 spike cleavage by the host proteases furin and TMPRSS2. We provide direct evidence by MS-glycoproteomics that identifies GalNAc-T1 as the glycosyltransferase initiating Thr678 glycosylation in the living cell. We demonstrate that the presence of elaborated glycans on Thr678 reduce proteolytic cleavage by TMPRSS2 and that a negative charge (via sialic acid) on Thr678-containing glycopeptides completely abrogates furin activity. We further confirm that mutations on Pro681 (present in major VOCs Alpha, Delta and Omicron) impair glycosylation of Thr678 and may therefore promote proteolytic processing of spike. We firmly establish O-glycosylation as a major determinant of SARS-CoV-2 spike maturation and propose disruption of O-GalNAc glycosylation as a considerable evolutionary driver for the emergence of SARS-CoV-2 VOCs.

## RESULTS AND DISCUSSION

Establishing a protein as a GalNAc-T substrate classically features expression in cells either lacking or overexpressing the respective GalNAc-T, followed by detection by GalNAc-recognising lectins.^72,73,85–87,99^ While generally powerful, identifying the modified glycosylation sites is often challenging by these methods, due to the interplay and ensuing compensatory effects between GalNAc-T isoenzymes. Bump-and-hole engineering enables a direct relation to the engineered GalNAc-T isoenzyme by introduction of a GalNAc analogue which can be bioorthgonally tagged and detected by various analytical techniques (**Figure *2***). We incubated recombinant SARS-CoV-2 spike WT, P681R or P681H constructs produced in human Expi293F cells with recombinant WT- or bump-and-hole engineered GalNAc-T1 (BH = I238A/L295A double mutant) or T2 (BH = I253A/L310A double mutant) and the bumped nucleotide sugar UDP-GalN6yne.^93^ We then tagged the glycosylated peptides with biotin picolyl azide by Cu(I)-catalyzed azide-alkyne cycloaddition (CuAAC) and visualized glycosylation via streptavidin blot (**Figure 2a**). An intense, single band corresponding to the S1 fragment was observed when WT (Wuhan/WH04/2020) spike was incubated with BH-GalNAc-T1 (**Figure 2b**). Negligible signal was observed on preparations incubated with either BH-T2 or the corresponding WT-GalNAc-Ts. A single substitution of Pro681 to either His or Arg led to near-complete abrogation of glycosylation by BH-T1. These data suggest that recombinant WT-S1 contains a dedicated GalNAc-T1 glycosylation site which is absent in recombinant WT-FL-S and obstructed upon variant-specific mutation of Pro681.

**Figure 2.**
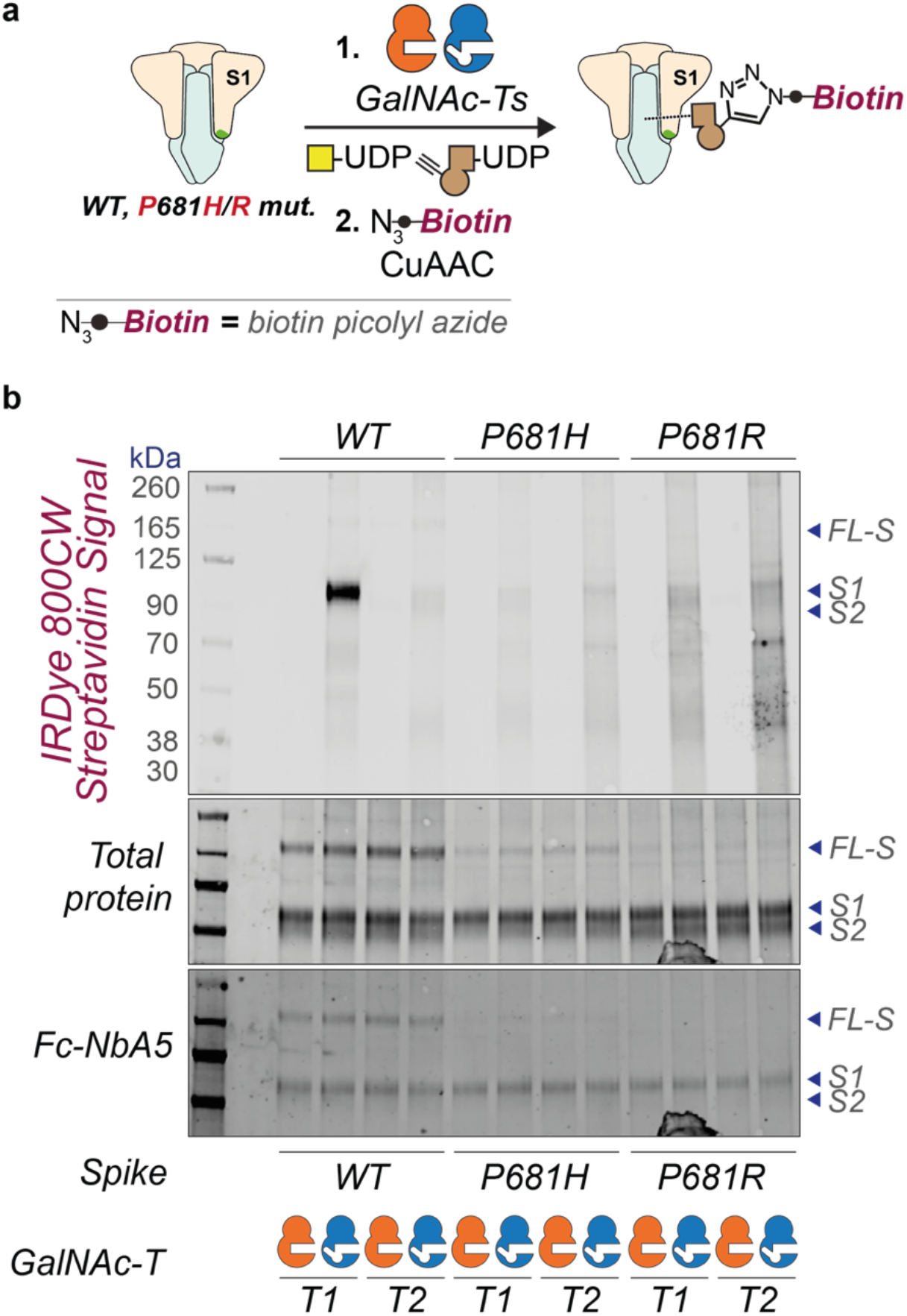
GalNAc-T isoenzyme-specific *in vitro* glycosylation of recombinant SARS-CoV2 spike preparations. **a)** Overview of chemoenzymatic experiments comparing WT and BH-GalNAc-T1/T2 glycosylation (and click-biotinylation) of WT and P681 mutant (P681H and P681R) spike preparations. **b)** Streptavidin blot of the chemoenzymatically tagged (glycosylated and biotin picolyl azide CuAAC-ligated) recombinant SARS-CoV-2 spike WT and P681 mutants. Visualised via IRDye 800CW-streptavidin fluorescence. FL-S: Full-length SARS-CoV-2 spike; S1: Cleaved SARS-CoV-2 spike S1 domain; S2: Cleaved SARS-CoV-2 spike S2 domain; Fc-NbA5: Fc-conjugated SARS-CoV-2 RBD spike-specific nanobody.

We then used a panel of synthetic peptides to study the effect of spike mutations on GalNAc-T1-mediated glycosylation. The peptide panel included variant-related mutations at the major hotspots: Gln675, Gln677, Asn679 and Pro681 (**Figure 3a**). GalNAc-T1 generated both mono- and di-glycopeptides from a WT (Wuhan) substrate peptide (**Figure 3b**). Consistent with the work by Ten Hagen and colleagues,^72^ notable reductions in glycosylation were observed for P681H and P681R peptides with approx. 80% and 90% of starting material remaining in the reaction mixture, respectively. These results validated the importance of a Pro in position +3 for GalNAc-T1 glycosylation.^100^ Single mutations at positions 675, 677, and 679, including N679K found in Omicron, reduced the amount of di-glycosylation but largely retained mono-glycosylation with almost complete consumption of the starting material. In contrast, the combination of two mutations at positions 675 and 677 showed a critical reduction of glycosylation. This was evidenced by reactions with the double mutant peptides Q675H+Q677H and Q675H+Q677R resulting in 9.7% and 5.7% conversion, respectively. These data confirmed the dependence of GalNAc-T1 on Pro681 but indicate that only certain VOC mutations impact O-GalNAc glycosylation. Mass spectrometry with electron-transfer dissociation (ETD) fragmentation revealed that mono-glycopeptides are exclusively GalNAc-modified on Thr678 while di-glycopeptides are modified at Thr676 and Thr678, indicating a hierarchy of sites where Thr678 is glycosylated first (**Figure 3c** and **Supporting Figure 1**). When BH-engineered GalNAc-T1 and UDP-GalN6yne were used in the *in vitro* assay with WT spike-derived peptides, we observed the same trends, confirming that BH-T1 recapitulates the substrate specificity of WT-T1 (**Figure 3c** and **Supporting Figure 1**).

**Figure 3.**
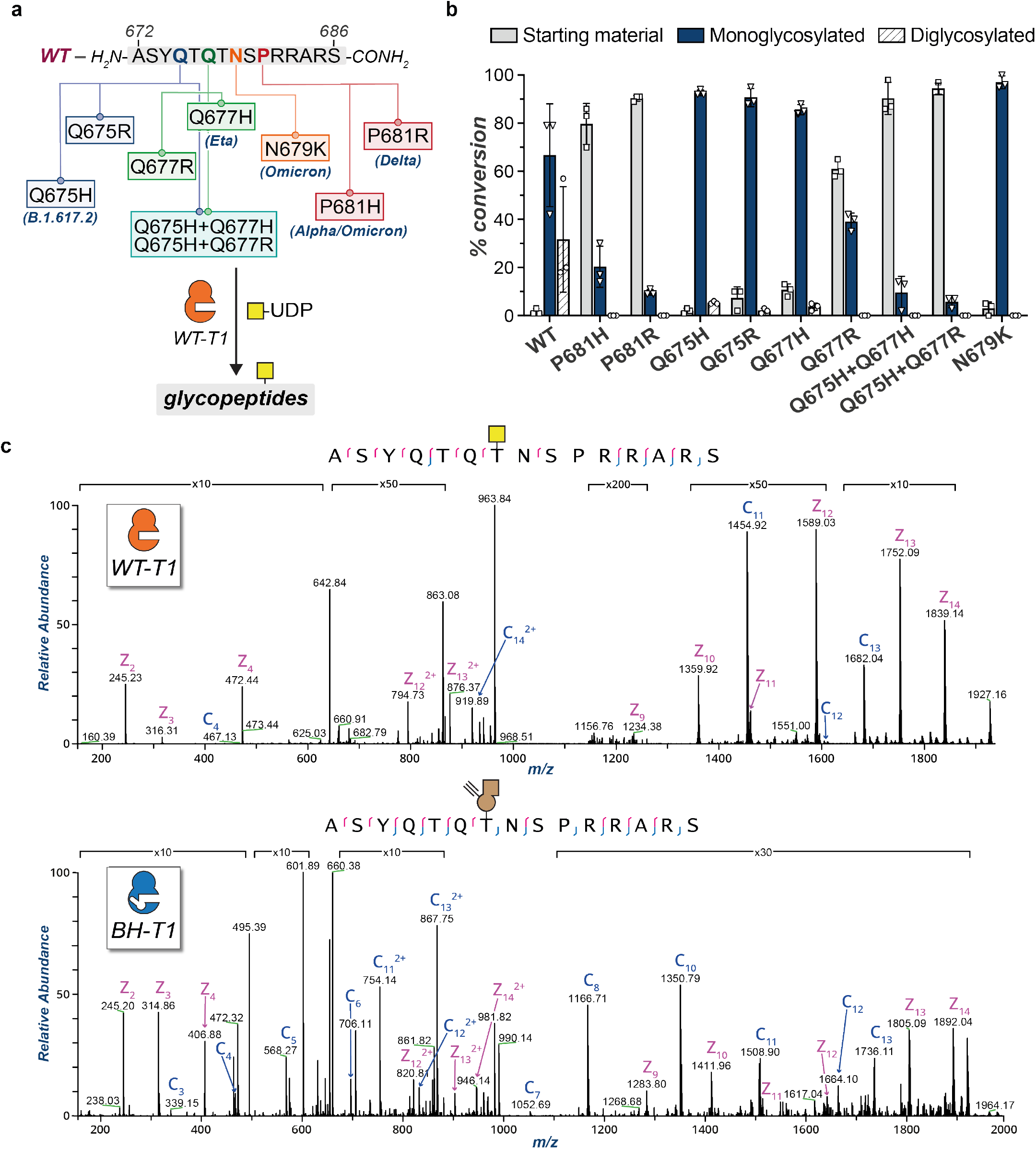
Evaluation of GalNAc-T1-mediated glycosylation on synthetic peptides. **a)** Peptide panel of the FCS proximal region in SARS-CoV-2 spike including WT and 9 mutant peptides. **b)** *In vitro* glycosylation results with recombinant WT-GalNAc-T1 and UDP-GalNAc assessed by LC-MS. Data are means ± SD from three independent replicates. **c)** Tandem MS (ETD) spectra for WT (*top*) and BH- (*bottom*) GalNAc-T1 glycosylation of the WT peptide H-ASYQTQTNSPRRARS-NH_2_ *in vitro*. Synthetic peptides were run on an Orbitrap Eclipse (Thermo) and subjected to ETD, followed by manual validation and hand curation. Legend: c ions are indicated in blue and z ions in pink. Yellow square = GalNAc. Brown alkyne = GalN6yne. WT-T1 = WT-GalNAc-T1. BH-T1 = BH-GalNAc-T1.

Having established a link between VOC mutations and glycosylation, we sought to rule out an immunological implication of the corresponding (glyco-)peptides that might impact any mechanistic deductions. Peptides WT, P681H and WT-GalNAc (**Figure 3a**; **“**WT-GalNAc” corresponds to the glycosylation product of the WT peptide carrying an O-GalNAc on Thr678), were evaluated in peripheral blood mononuclear cells (PBMC) from n=48 SARS-CoV-2 vaccinated individuals for T cell interferon-gamma (IFN-γ) secretion using an Enzyme-linked immunosorbent spot (ELISpot) assay.^101^ As shown in **Supporting Figure 2**, the median [IQR] response to the spike protein was 33.5 [16.7-69] spot forming cells (SFC) per million PBMC and to the combined pool of M and N proteins was 10 [0-29] SFC/million PBMC. The median [IQR] response to the peptide WT-GalNAc was 0 [0-3.7] in n=44 individuals, compared to the peptide P681H 0 [0-3.5] in n=21, and WT 10 [0-27] in n=3 individuals. These results indicate that neither (glyco-)peptide is a T cell target in vaccinated individuals.

### GalNAc-T selective MS-glycoproteomics analysis allows O-glycosite and glycan composition investigation *in vitro* and in engineered cells

Assigning the activity of GalNAc-T isoenzymes to specific glycosylation sites is complicated by the redundant nature of GalNAc-Ts and the complex dynamics of the secretory pathway, neither of which can be accurately replicated with *in vitro* assays using synthetic peptides. Furthermore, the FCS-adjacent region lacks cleavage sites of the proteases most commonly used in MS sample preparation, resulting in large glycopeptides further hampering analysis. The use of specialized chemical tools can address these shortcomings and report on GalNAc-T activity in the secretory pathway of living cells while offering a bioorthogonal handle to aid MS analysis. We stably transfected Expi293F cells with constructs for both SARS-CoV-2 spike (Wuhan) and either WT- or BH-versions of GalNAc-T1 or T2, along with the biosynthetic machinery to generate UDP-GalN6yne in the cell from a membrane permeable precursor (Ac_4_GalN6yne) that was introduced via cell feeding (**Figure 4a**).^98^ Spike samples were isolated and derivatized by CuAAC with an azide-functionalized imidazolium group containing a permanent positive charge (ITag-azide).^97,102–104^ This treatment introduced GalN6yne in an isoenzyme-specific fashion while endowing glycopeptides with an additional positive charge that facilitates MS-analysis.^97^ The separated FL-S and S1/S2 fractions were subjected to in-gel digestion and analyzed by tandem MS. We used high intensity collision-induced dissociation (HCD) to obtain naked peptide sequences and glycan compositions, and then used the ITag-containing GalN6yne diagnostic ion to trigger ETD fragmentation of the peptide backbone (**Figure 4b**).^71,97,105,106^ Through both computational (Byonic, ProteinMetrics) and manual validation, we found that Thr678 carried ITag-modified GalN6yne in both FL-S and S1 samples exclusively in cells expressing BH-T1, but not BH-T2 or any WT-GalNAc-Ts (**Figure 4c** and **Supplementary Data 1**). The additional positive charge of the ligated ITag permitted straightforward ETD fragmentation of a 21-amino acid glycopeptide. In contrast, the corresponding glycopeptide in samples expressing WT-T1 could not be unambiguously sequenced, highlighting the ability of chemical tools to help advance site-specific O-glycoproteomics. We further found that both BH-T1 and BH-T2 glycosylated Thr323, a previously detected glycosylation site that had thus far not been associated with any GalNAc-T isoenzyme (**Figure 4c** and **Supporting Figure 3**). These results were also recapitulated *in vitro* through glycosylation of recombinantly expressed spike with recombinantly expressed soluble constructs of BH-GalNAc-T1 and BH-GalNAc-T2, followed by CuAAC ligation of ITag-azide and MS-glycoproteomics analysis (SI and **Supporting Figure 4**). Our data directly proved that GalNAc-T1 glycosylates Thr678 in the living cell.

**Figure 4.**
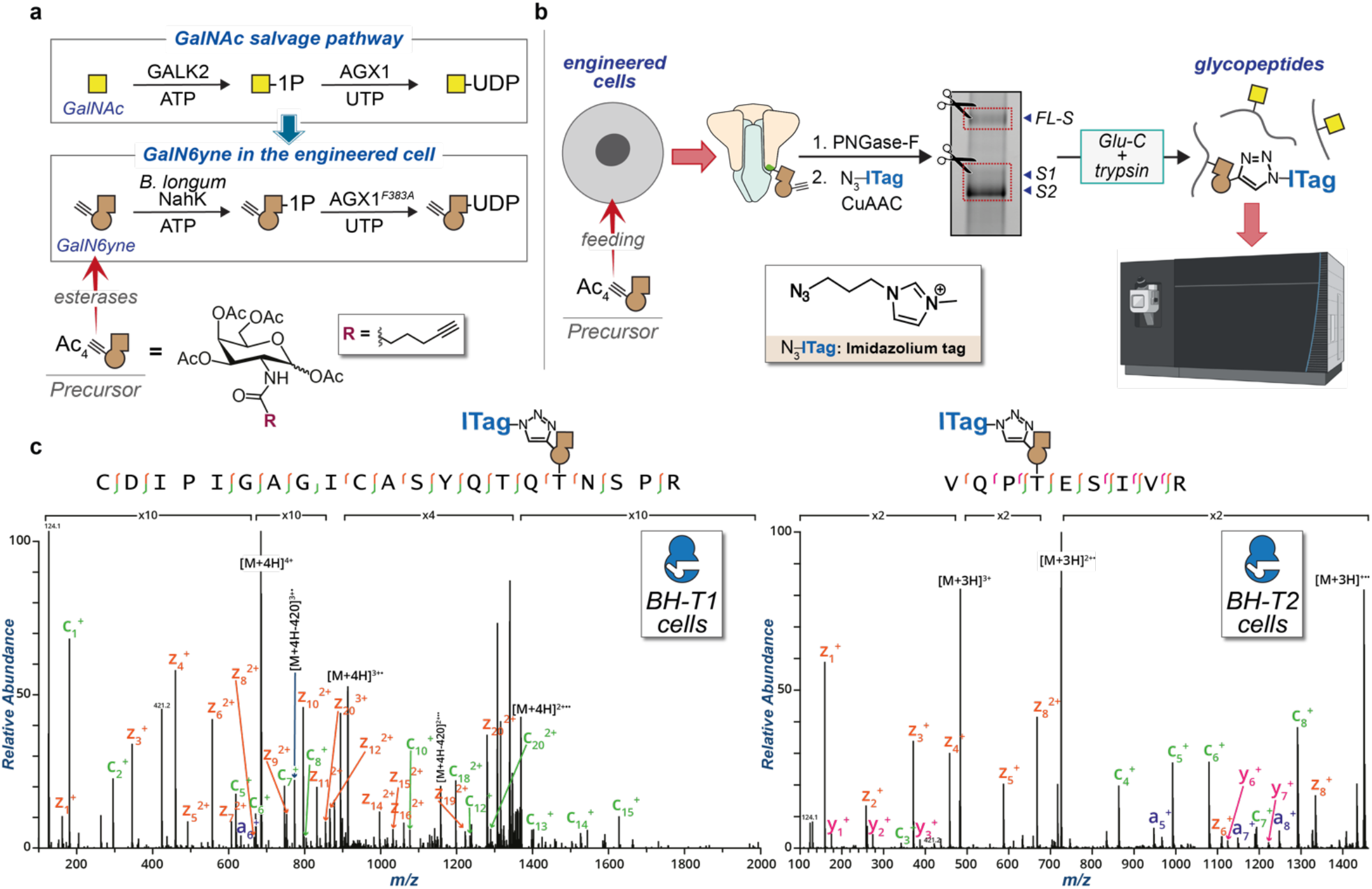
Uncovering the relationship between GalNAc-T1 and Thr678 by chemical tools. **a)** GalNAc salvage pathway and UDP-GalN6yne biosynthesis. Expression of the kinase NahK and the pyrophosphorylase AGX1^F383A^ permits biosynthesis of UDP-GalN6yne in engineered cells. **b)** Graphical representation of the MS-glycoproteomics methodology for engineered cells: SARS-CoV-2 spike was recombinantly expressed in Expi293F cells co-expressing NahK, AGX1^F383A^ and either BH-GalNAc-T1 or T2. Following isolation and de-*N*-glycosylation, ITag-azide was introduced by CuAAC and the protein preparation subjected to in-gel digestion and MS by HCD-triggered ETD. **c)** Annotated tandem MS (ETD) spectra of the major hits tagged by BH-GalNAc-T1 (*left*) and BH-GalNAc-T2 (*right*). c ions are indicated in green, z ions in orange and y ions in pink. Yellow square = GalNAc. Brown alkyne = GalN6yne. Ac_4_GalN6yne precursor = membrane permeable peracetylated GalN6yne. BH-T1 cells = Expi293F cells co-transfected with WT SARS-CoV-2 spike and BH-GalNAc-T1. BH-T2 cells = Expi293F cells co-transfected with WT SARS-CoV-2 spike and BH-GalNAc-T2.

### Elaborated O-linked sialoglycans on Thr678 confer proteolytic resistance to SARS-CoV-2 spike

Glycosylation has the potential to modulate the proteolytic processing of a peptide depending on the distance to the cleavage site and glycan composition.^88,91,92^ We used a direct method to investigate whether O-GalNAc glycans on Thr678 modulate cleavage by furin. To this end, we designed synthetic Förster Resonance Energy Transfer (FRET)-active substrate peptides to assess proteolytic activity. Peptides spanning residues 672 to 689 contained N-terminal 2-aminobenzoyl (Abz) and C-terminal 3-nitrotyrosine (3-NO_2_Tyr) as fluorescence donor and quencher moieties, respectively. An increase in fluorescence intensity indicated proteolytic cleavage (**Figure 5a**).^107^ We first compared non-glycosylated substrates corresponding to either WT (**FRET-1**) and P681H mutant spike (**FRET-2**). The P681H mutation had no discernable effect on the rate of furin-mediated cleavage, confirming the data by Whittaker and colleagues that the addition of a basic amino acid is not by itself a defining characteristic of spike furin cleavage enhancement in existing VOCs.^55^

**Figure 5.**
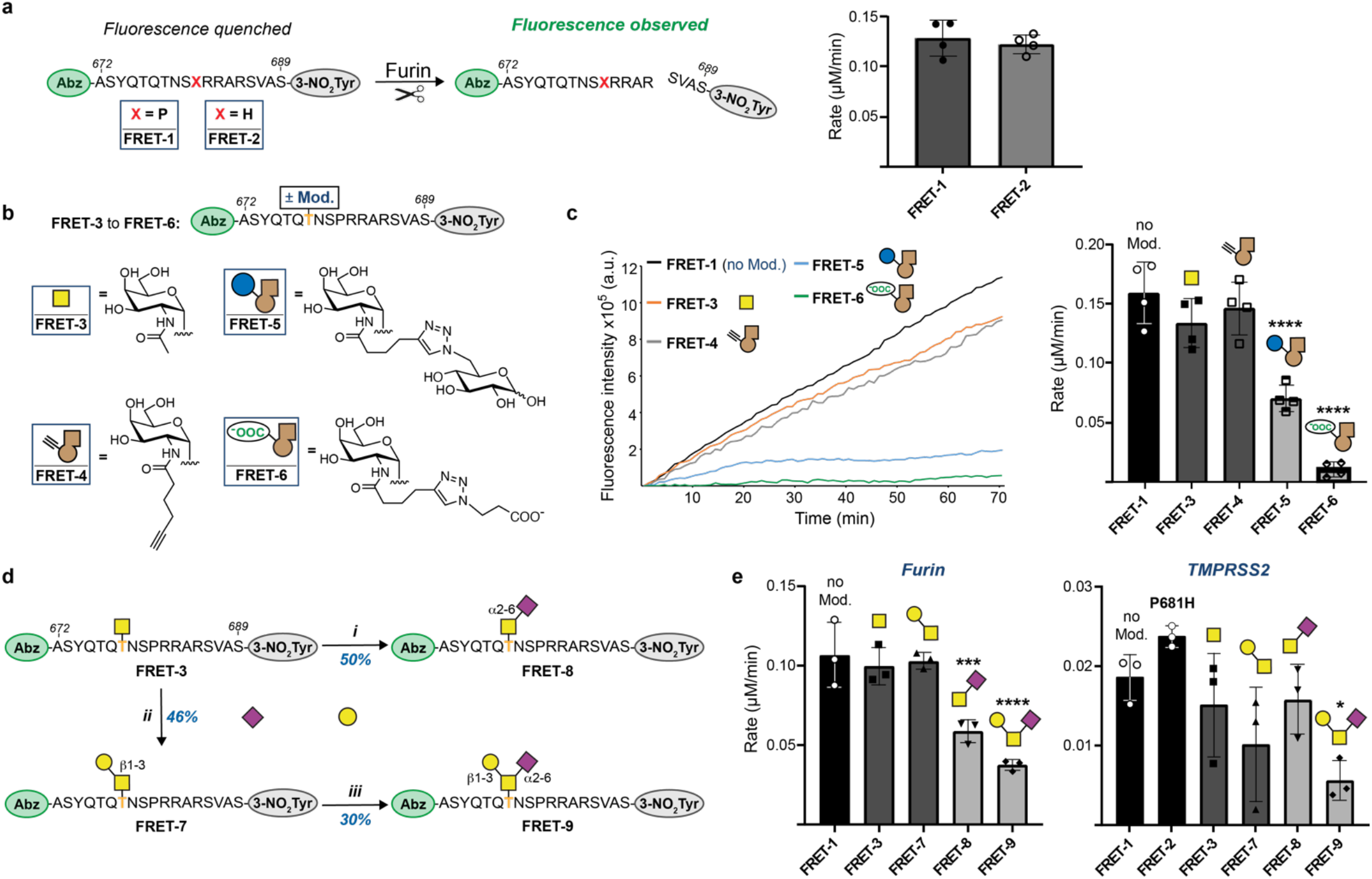
Chemical elaboration of O-glycosylation to assess proteolytic cleavage of glycopeptides. **a)** Experimental design and comparison of furin cleavage between WT and P681H peptide substrates. Peptides containing N-terminal 2-aminobenzoate (Abz) as a FRET donor, and C-terminal 3-nitrotyrosine (3-NO_2_Tyr) as a FRET quencher that was removed upon proteolytic cleavage. **b)** Chemical modifications of synthetic (glyco-)peptides **FRET-3** through **FRET-6**, generated via *in vitro* glycosylation and CuAAC. **c)** *Left*: Time course of fluorescence increase upon furin cleavage reactions of 20 µM **FRET-1** and **FRET-3** to **FRET-6** with 0.8 U/mL furin. Linear fluorescence increase is shown and normalised to the corresponding control run without furin. *Right*: Rates of furin cleavage of glycopeptides **FRET-1** and **FRET-3** to **FRET-6** obtained through linear regression and normalisation to control runs without furin. Data are means ± SD of four independent experiments. **d)** Chemoenzymatic synthesis of glycopeptides **FRET-7** through **FRET-9:** *i*. ST6GalNAc-1 (150 µg/mL), CMP-Neu5Ac (1.5 eq.), pH 7.5, 37°C for 16 hours, 46% yield; *ii. Dm*C1GalT1 (1 µM), UDP-Gal (1.5 eq.), pH 7.5, 37°C for 16 hours, 50% yield; *iii*. ST6GALNAC-2 (10 µg/mL), CMP-Neu5Ac (1.5 eq.), pH 7.5, 37°C for 48 hours, 30% yield. Bold and orange T denotes Thr678 in the glycopeptides. **e)** Rates of furin (*left*) and TMPRSS (*right*) cleavage of (glyco-)peptides **FRET-1** to **FRET-3** and **FRET-7** to **FRET-9** obtained through linear regression and normalization to control runs without furin. Data are means ± SD of three independent experiments. Group comparison was performed via one-way ANOVA with Tukey’s multiple comparisons test and asterisks annotate P values: \**P* < 0.0332 ; \*\**P* < 0.0021; \*\*\**P* < 0.0002; \*\*\*\**P* < 0.0001, compared to the non-modified peptide (**FRET-1**). CMP = cytidine monophosphate; CIAP = Calf intestinal alkaline phosphatase. Neu5Ac = N-acetylneuraminic acid.

We hypothesized that an increase of furin processing in mutant spike may not stem directly from recognition of the bare peptide sequence, but rather a decreased capacity of GalNAc-T1 to introduce O-GalNAc glycans to peptides with mutations proximal to the recognition site. We thus tested whether glycosylation of furin substrate peptides impacts proteolytic cleavage. FRET reporter peptides carrying GalNAc (**FRET-3**) or alkyne-containing GalN6yne (**FRET-4**) were synthesized by chemoenzymatic glycosylation with WT- and BH-GalNAc-T1, respectively. Glycosylation with the single monosaccharides alone did not substantially impact the furin cleavage rate compared to the WT peptide **FRET-1** (**Figure 5c** and **Supporting Figure 5**). We speculated that elaboration of GalNAc to larger or charged glycans might introduce additional structural constraints on furin recognition. The alkyne tag present on GalN6yne gave an opportunity to modify the biophysical properties of glycopeptides in a straightforward fashion, enabling synthetic efforts to furnish glycopeptides with specific additional groups or functionalities. We reacted glycopeptide **FRET-4** with two organic azides under CuAAC conditions: 6-azido-6-deoxy-glucose yielded the pseudodisaccharide **FRET-5**, while 3-azido-propionic acid introduced an additional acidic functionality to investigate the impact of a negative charge on furin cleavage in glycopeptide **FRET-6**. Both click-elaborated glycopeptides displayed a significant reduction in furin cleavage (**Figure 5c** and **Supporting Figure 5**). **FRET-5** exhibited an 80% decrease in the rate of furin cleavage with respect to **FRET-1**, which is attributable to the relative steric expansion. Strikingly, **FRET-6** which carried a smaller, negatively charged modification, resulted in a 93% rate reduction, almost completely abrogating furin activity. We concluded that the elaboration of O-glycans on Thr678, especially with negatively charged modifications, severely impedes the activity of furin.

Sialic acid, a common capping monosaccharide of O-glycans, is negatively charged under physiological pH. We reasoned that the presence of sialic acid might modulate furin cleavage, and hence chemoenzymatically synthesized a set of novel, spike-derived glycopeptides to test on our cleavage assay. First, *Drosophila melanogaster* C1GALT1 was used to extend **FRET-3** with β3-linked galactose to give **FRET-7**.^108^ α2,6-Linked sialic acid was introduced into **FRET-3** and **FRET-7** using the enzymes ST6GALNAC1 and ST6GALNAC2, respectively, to yield the sialoglycopeptides **FRET-8** and **FRET-9** (**Figure 5d**).^109^ While the uncharged disaccharide in **FRET-7** only had a marginal (3.5% decrease) effect on furin rate compared to the parental peptide **FRET-1**, the presence of a sialic acid led to a striking 45% reduction of furin rate in glycopeptide **FRET-8** and a 65% reduction in glycopeptide **FRET-9** (**Figure 5e**). We concluded that the elaboration of O-glycans on Thr678 with negatively charged sialic acid residues severely hampers furin activity.

To our knowledge, in contrast to furin,^88^ TMPRSS2 has not been comprehensively probed for cleavage of glycopeptide substrates. We sought to establish whether O-GalNAc glycans could modulate TMPRSS2 activity in a similar fashion to furin. Recombinant TMPRSS2 was subjected to our selection of (glyco-)peptide FRET substrates **FRET-1, FRET-3** and **FRET-7** to **FRET-9** (**Figure 5e**). While all glycans somewhat impacted TMPRSS2 activity with respect to the non-glycosylated peptide **FRET-1**, the trisaccharide-containing sialoglycopeptide **FRET-9** (70% rate reduction) impacted cleavage more drastically than all other (glyco-)peptides.

Our *in vitro* glycosylation experiments suggested that S1 is the only available GalNAc-T1 substrate on WT spike after secretion from human cell culture (**Figure 2b**). Such behaviour could be explained by the presence of elaborated, sialylated O-glycans on recombinant FL-S, which would both prevent furin cleavage and block access by GalNAc-T1 *in vitro*. This would suggest an enrichment of sialylated O-glycans on FL-S relative to processed S1. To explore this notion, we subjected the FL-S and S1/S2 gel bands from a recombinant WT-spike preparation to MS-glycoproteomics analysis, searching for both simple and elaborated O-GalNAc glycans. By calculating the intact masses of various expected O-glycopeptides in recombinant spike and then obtaining the associated extracted ion chromatograms (XICs), we found that over 5-fold higher glycopeptide signal is present in FL-S when compared to cleaved S1/S2 fractions (**Figure 6** and **Supplementary Data 2**). Furthermore, when accounting for sialic acid-containing glycopeptides only, an ∼8-fold higher abundance was observed for FL-S relative to S1/S2 (**Supplementary Data 2)**.

**Figure 6.**
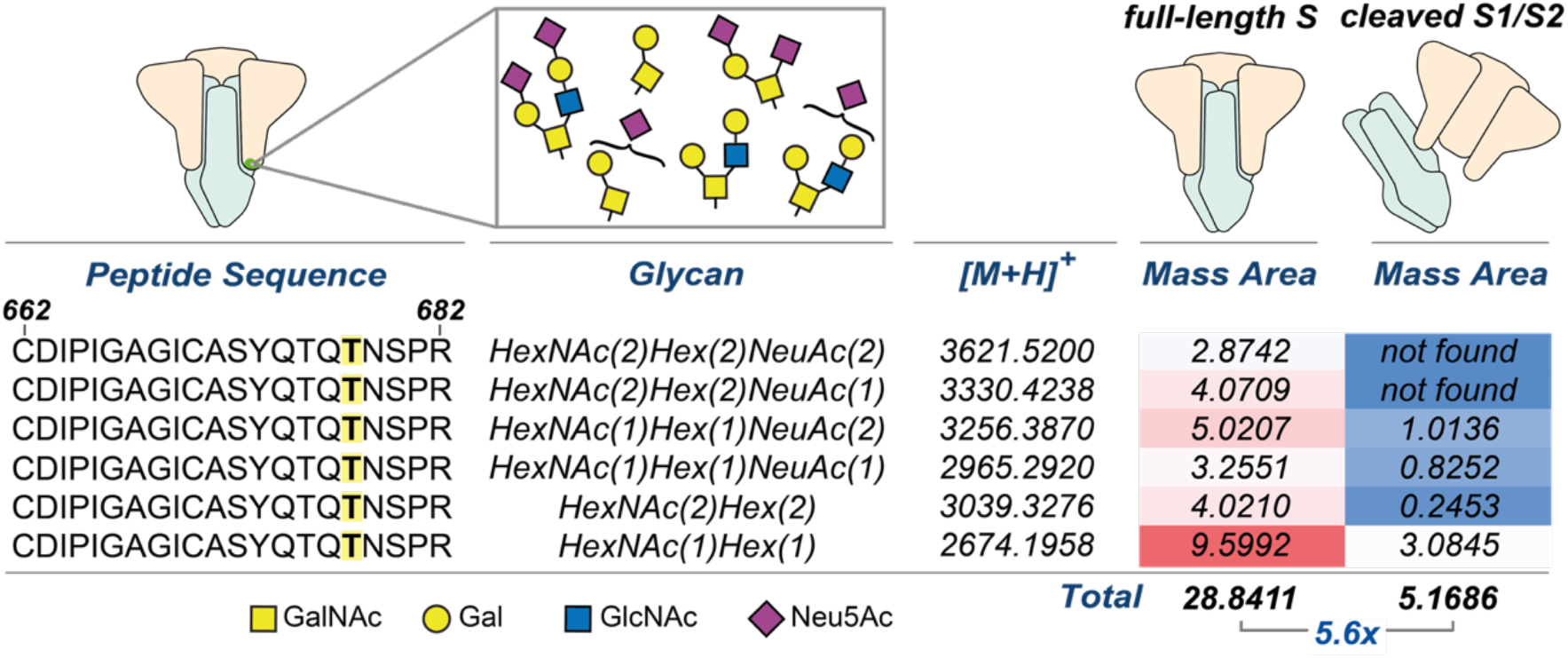
Extracted ion abundance for O-glycopeptides from FL-S or S1/S2 recombinant spike fractions. Mass areas were normalised against their corresponding base peaks and multiplied by a factor of 10,000. All data analysis was performed in Thermo XCalibur software and hand-curated. Highlighted T corresponds to Thr678 which was found to be modified by diverse O-glycans.

## DISCUSSION

Our data strongly indicate that elaborated, negatively charged O-GalNAc glycans on Thr678 of SARS-CoV-2 spike have a negative effect on proteolytic cleavage. Such glycans are produced on lung epithelial cells which express GalNAc-T1,^72^ suggesting that glycosylation is a physiologically relevant modification that could restrict the maturation (by proteolysis) of spike in WT SARS-CoV-2.^110^ The propensity of SARS-CoV-2 variants of concern to outcompete each other has been linked to both increased infectivity and immune escape. Within the evolutionary trajectory from the Alpha to Delta and Omicron variants, notable changes in the amino acid sequence proximal to the FCS indicated that proteolytic cleavage is gradually enhanced, congruent with their increased infectivity. Mutations of Pro681 have been detected in early variants such as Alpha (P681H). We found that this mutation did not increase the rate of cleavage by both furin and TMPRSS2, further implicating glycosylation as a restricting factor for spike processing. Notably, the analogous mutation found on the more transmissible Delta variant (P681R) has been linked to an increase in furin cleavage by Whittaker and colleagues,^43^ suggesting an evolutionary trajectory that places suppression of O-glycosylation before increasing intrinsic furin recognition. This is further underlined by the mutations found in Omicron, which features both the P681H with the N679K mutations: in contrast to P681H and consistent with mapped amino acid preferences of GalNAc-T1,^111^ the N679K mutation does not substantially impact glycosylation, but it leads to enhanced furin cleavage of synthetic peptides.^43^ These FCS-adjacent mutations therefore act synergistically and have likely evolved to both suppress glycosylation and intrinsically enhance furin cleavage. Since the closest relatives to SARS-CoV-2, the strains RaTG13 Bat-CoV and GD Pangolin-CoV, share the exact same peptide sequence without harbouring an FCS (**Figure 1**), we speculate that O-glycosylation might have been a remnant of ancestral strains that has been subsequently lost through the process of viral evolution.

The presence of O-GalNAc glycans adjacent to proteolytic cleavage sites has been found to impact processing of secreted proteins.^88–92^ The glycosylation site at Thr678 of spike is not in direct proximity of the FCS, potentially explaining why a single GalNAc residue is not sufficient to modulate furin activity and only minimally impacts TMPRSS2 activity. The necessity for the glycan to be elaborated or sialylated to reveal the negative effect upon the rates of cleavage further highlights the need for accurate glycan tracing techniques._112_ Tuning the chemical properties of glycopeptides in a straightforward fashion by CuAAC was pivotal in enabling an initial understanding of the substrate-activity relationship of proteases. This strategy informed the targeted synthesis of elaborated O-glycopeptide substrates, providing a convenient method to fine-tune substrate scope in a time- and resource-efficient manner to address specific hypotheses. Chemical tools thus provided insights into glycosyltransferase specificity, improved the efficacy of detection by detectability by MS, and allowed the exploration of protease substrate specificity, showcasing the power of such tools in biomedical discovery.

## ETHICS STATEMENTS

The Legacy study was approved by London Camden and Kings Cross Health Research Authority (HRA) Research and Ethics committee (REC) IRAS number 286469 and sponsored by University College London.

## ACKNOWLEDGMENTS

We thank David Briggs, Sarah Maslen, Neil McDonald, Phil Walker, Steve Gamblin and Nicola O’Reilly for valuable support and Theresa Zeisner, Jennifer Milligan and John Diffley for helpful discussions. We thank Kelly Ten Hagen and Nadine Samara for providing crucial insight. We thank Junwon Choi and Carolyn R. Bertozzi for UDP-GalN6yne. This work was supported by the Francis Crick Institute which receives its core funding from Cancer Research UK (FC001749 and FC0010219), the UK Medical Research Council (FC001749 and FC0010219), and the Wellcome Trust (FC001749 and FC0010219). This work was funded by the UK Biotechnology and Biological Sciences Research Council (BB/V008439/1 to E. G.-R., L. D. V. and B. S., BB/V014862/1 to G. B.-T. and B. S. and BB/M028836/1 to S. L. F.), the Engineering and Physical Sciences Research Council (EP/S005226/1 to S. L. F.), the Wellcome Trust (218304/Z/19/Z to A. M. and B. S.) and the European Research Council (788231 to S. L. F.). M. Z.-H. was supported by a Crick-HEI studentship funded by the Department of Chemistry at Imperial College London and the Francis Crick Institute. S. A. M. is supported by the Yale Science Development Fund and NIGMS R35 GM147039. K. E. M. is supported by a Yale Endowed Postdoctoral Fellowship. R.J.W. receives support from Rosetrees (M926). R.H.-G., thanks ARAID, the Agencia Estatal de Investigación (AEI, BFU2016-75633-P and PID2019-105451GB-I00) and Gobierno de Aragón (E34_R17 and LMP58_18 to R.H.-G.) with FEDER (2014–2020) funds for “Building Europe from Aragón” for financial support. The PBMC portion of this work was supported jointly by the National Institute for Health Research (NIHR) University College London Hospitals Department of Health’s NIHR Biomedical Research Centre (BRC) and core funding from the Francis Crick Institute, which receives its funding from Cancer Research UK, the UK Medical Research Council, and the Wellcome Trust. EW is supported by the BRC’s funding scheme. For the purposes of Open access, the author has applied a CC-BY copyright to any author accepted version of a manuscript arising from this submission.

