## supplementary information for "O-Linked Sialoglycans Modulate the Proteolysis of SARS-CoV-2 Spike and Likely Contribute to the Mutational Trajectory in Variants of Concern"

[a]Chemical Glycobiology Laboratory, The Francis Crick Institute, NW1 1AT London, United Kingdom. [b]Department of Chemistry, Imperial College London, W12 0BZ London, United Kingdom. [c]Signalling and Structural Biology Lab, The Francis Crick Institute, London, NW1 1AT, United Kingdom. [d]Proteomics Science Technology Platform, The Francis Crick Institute, NW1 1AT London, United Kingdom. [e]Department of Chemistry, Yale University, 275 Prospect Street, New Haven, CT, 06511, USA. [f]Structural Biology Science Technology Platform, The Francis Crick Institute, NW1 1AT London, United Kingdom. [g]Structural Biology of Disease Processes Laboratory, Francis Crick Institute, London NW1 1AT, United Kingdom. [h]Manchester Institute of Biotechnology, University of Manchester, 131 Princess Street, Manchester M1 7DN. [i]Chemical Biology Science Technology Platform, The Francis Crick Institute, London NW1 1AT, UK. [j]Institute of Biocomputation and Physics of Complex Systems, University of Zaragoza, Zaragoza, Spain. [k]Copenhagen Center for Glycomics, Department of Cellular and Molecular Medicine, University of Copenhagen, Copenhagen, Denmark. [l]Fundación ARAID, 50018 Zaragoza, Spain. [m]Tuberculosis Laboratory, The Francis Crick Institute, London, UK. [n]Wellcome Centre for Infectious Diseases Research in Africa, University of Cape Town, Observatory, South Africa. [o]Department of Infectious Diseases, Imperial College London, London, W12 0NN UK. [p]Institute of Infectious Disease and Molecular Medicine and Department of Medicine, University of Cape Town, Observatory, South Africa. [q]The Francis Crick Institute, London NW1 1AT, UK. [r]University College London Hospitals (UCLH) Biomedical Research Centre, London, UK.

*

### SUPPORTING FIGURES

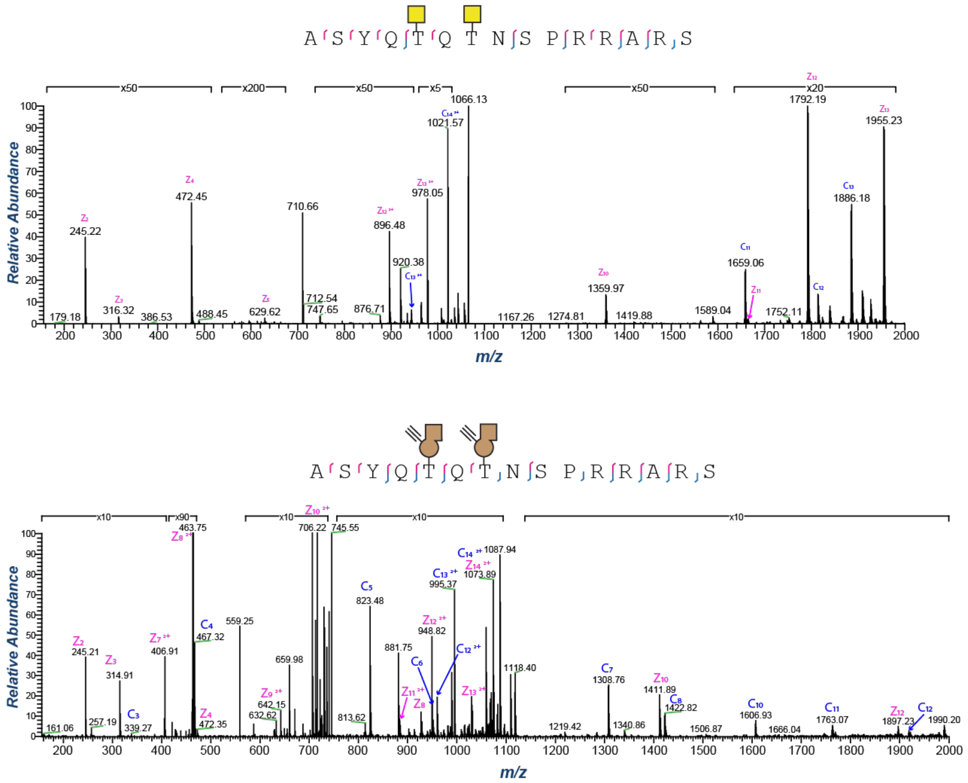

**Supporting Figure 1:** In vitro synthetic peptide double glycosylation results with WT-GalNAc-T1 and UDP-GalNAc (*top*) or BH-GalNAc-T1 and UDP-GalN6yne (*bottom*) assessed by tandem MS (ETD). Legend: c ions are indicated in blue and z ions in pink.

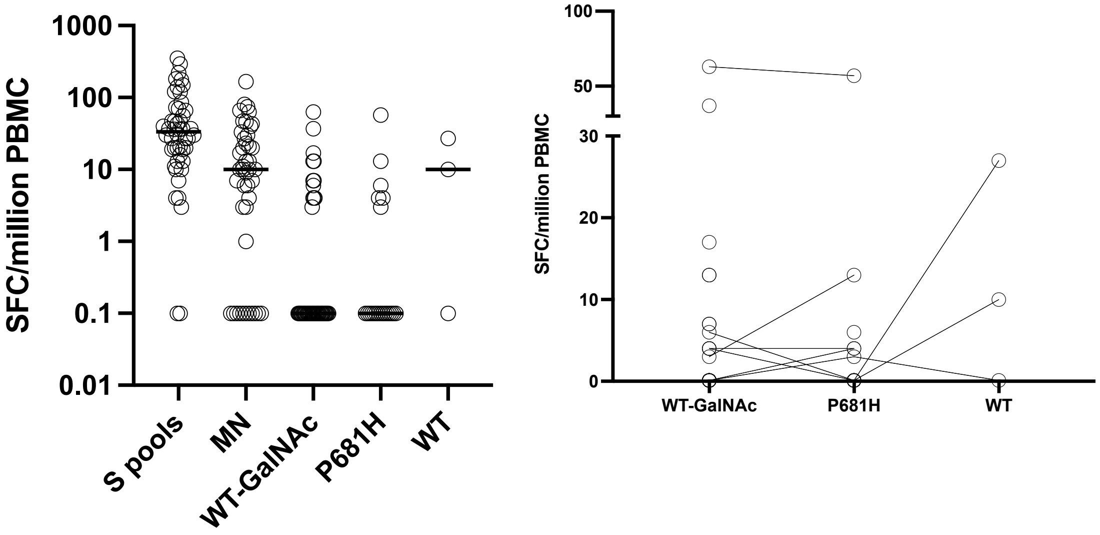

**Supporting Figure 2:** Interferon-gamma ELIspot results expressed as Interferon-gamma Spot Forming Cells per million PBMC, following correction for the unstimulated background by subtraction. Responses to peptide pools covering S, MN (used as a combined pool), as well as individual peptides wild type (WT), P681H and WT-GalNAc are shown. Tested in a range of vaccinated individuals (n=48), the median [IQR] response to the Spike protein was 33.5 [16.7-69] and to the combined pool of M and N proteins was 10 [0-29] SFC/million PBMC. The median [IQR] response to the peptide WT-GalNAc was 0 [0-3.7] in n=44 individuals, compared to the peptide P681H 0 [0-3.5] in n=21, and WT 10 [0-27] in n=3 individuals.

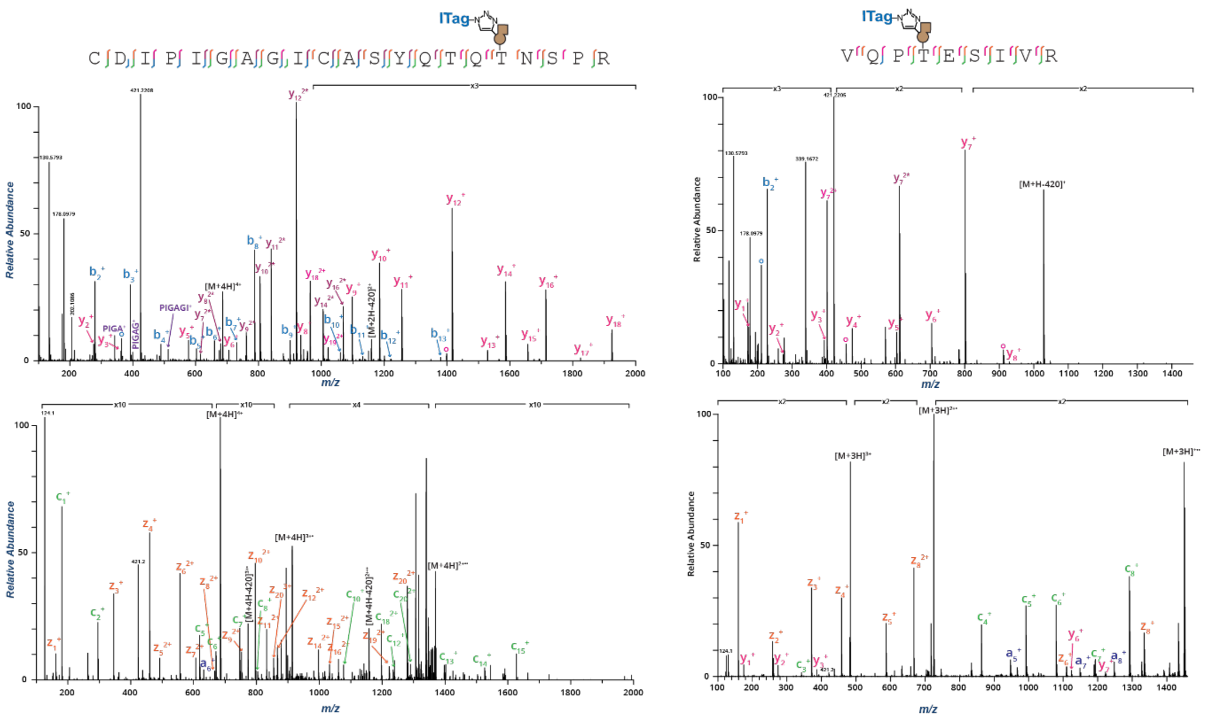

**Supporting Figure 3:** Annotated tandem MS (top, HCD; bottom, ETD) spectra of the major hits tagged by BH-GalNAc-T1 (*left*) and BH-GalNAc-T2 (*right*) from engineered cells co-expressing WT SARS-CoV-2 spike and BH-GalNAc-T1 or BH-GalNAc-T2. Legend: b ions are indicated in blue, y ions in pink, c ions in green and z ions in orange.

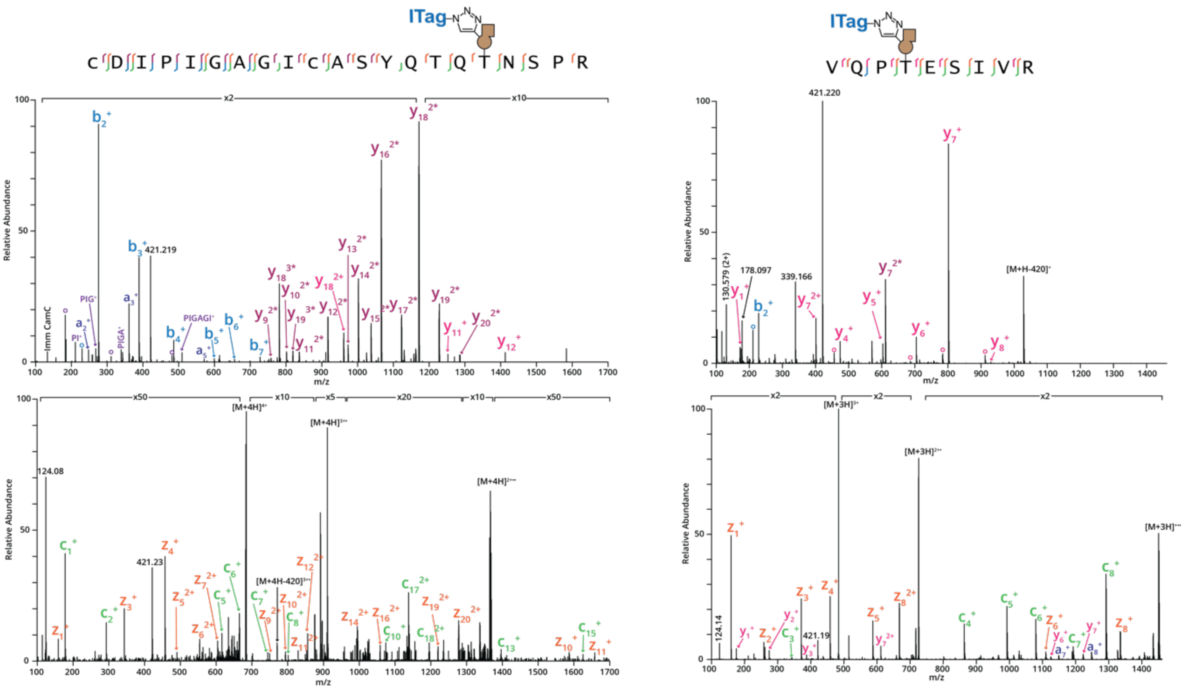

**Supporting Figure 4:** Annotated tandem MS (*top*, HCD; *bottom*, ETD) spectra of the major hits tagged by BH-GalNAc-T1 (*left*) and BH-GalNAc-T2 (*right*) from recombinant WT SARS-CoV-2 spike that was *in vitro* glycosylated with soluble BH-GalNAc-T1 or BH-GalNAc-T2. Legend: b ions are indicated in blue, y ions in pink, c ions in green and z ions in orange.

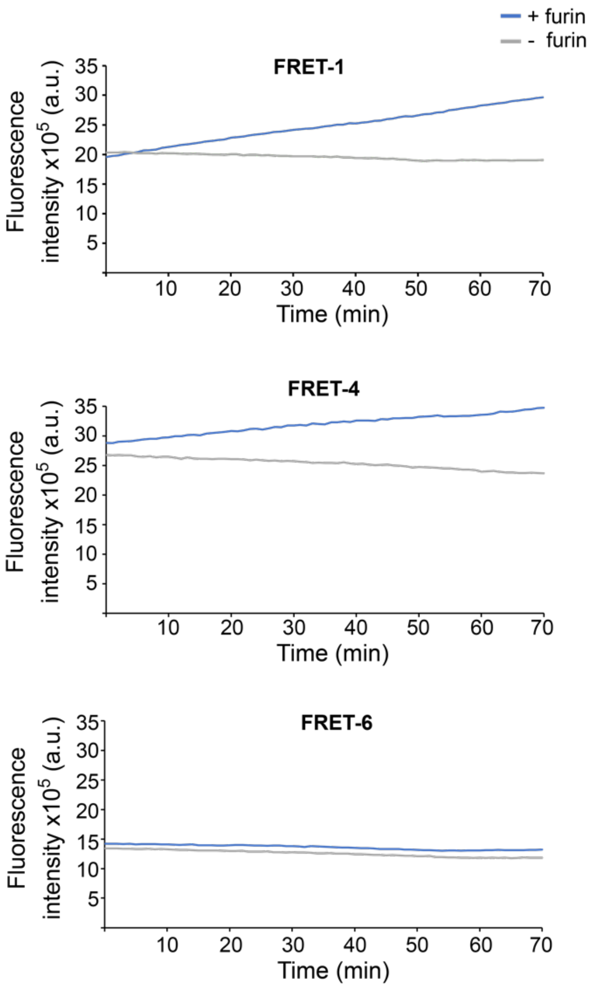

**Supporting Figure 5:** Fluorescence traces of furin cleavage experiments with (glyco-)peptides **FRET-1** and **FRET-3** to **FRET-6**. Time course of linear fluorescence increase is shown. Reactions contained 20 µM (glyco-)peptide and 0.8 U/mL furin or buffer as control. Data are from one representative of four independent replicates.

### MATERIALS AND METHODS

Unless otherwise stated, reagents were purchased from commercial sources and used without purification.

#### Protein expression and purification

FLAG-tagged constructs corresponding to the luminal domain of GalNAc-T1 WT or I238A/L295A ("BH") were expressed as described previously.[1] His-tagged constructs of luminal GalNAc-T2 WT or I253A/L310A (BH) were expressed as described before.[2] Photobacterium sp. JT-ISH-224 2,6-sialyltransferase (2,6-ST) was a gift from Sabine Flitsch.

Spike constructs for *in vitro* glycosylation were synthesized and cloned into pcDNA 3.1(+) by Genscript (Rijswijk, Netherlands) as described before[3,4] in pre-fusion stabilised "2P" (K986P and V987P) form:[5] Wuhan variant spike ectodomain (YP_009724390.1, residues 1-1208) and its versions with substitutions P681R or P681H.

The SARS-CoV-2 spikes were made as C-terminal His-tagged fusion proteins expressed and purified as described before.[3,4] In brief, proteins were expressed in Expi293F cells (Thermo Fisher, Waltham, USA) cultured according to the manufacturer's instructions in 8% $CO_2$ humidified atmosphere at 37 °C with shaking at 125 rpm, and transfected at $3\times10^6$ cells/mL density with ExpiFectamine 293 (Thermo Fisher) with 1 mg of DNA per 1 L of culture. The cells were transferred to a 32 °C incubator 24 h after the transfection and the supernatant harvested on the fourth day after the transfection[6]. The proteins were purified from the supernatant by affinity purification with HisPur Cobalt Resin (Thermo Fisher) followed by size exclusion chromatography with a Superdex 200 Increase 10/300 GL (GE Life Sciences, Piscataway, USA) column into a 20 mM Tris pH 8.0, 150 mM NaCl buffer.

SARS-CoV-2 spikes from BH-engineered cells were expressed in Expi293F (Thermo Fisher) cells. 30 mL cultures were grown in Freestyle 293 Expression Medium (Thermo Fisher) in an 8% $CO_2$ humidified atmosphere at 37 °C with shaking at 125 rpm and transfected with 1 mg/L of culture DNA with ExpiFectamine 293 (Thermo Fisher) according to the manufacturer's protocol. Transfection of the SARS-CoV-2 S protein expression constructs was performed first, followed by selection with geneticin (150 µg/mL) until cells returned to high viability (>95%) and normal growth rate. Then, cells were transfected with pSBbi vectors encoding full-length GalNAc-T, AGX1$^{F383A}$ and *B. longum* NahK as described previously[7], followed by selection with hygromycin B (150 µg/mL). For protein expression, cultures were expanded to a total media volume of 100 mL and transferred to a 32ºC incubator with 8% CO2 humidified atmosphere with shaking at 125 rpm. After passage and at +24 h, +48 h and +72 h, the media was either supplemented with $Ac_4$GalN6yne added to a final concentration of 50 µM (from a 50 mM $Ac_4$GalN6yne stock solution in DMSO), or the same volume of DMSO only was added. At +4 days, conditioned media was collected, clarified, buffer stocks solutions were added to a final concentration of 20 mM Tris pH 7.8 and 10 mM imidazole, and then incubated with 2×250 µL Ni-NTA agarose resin (Qiagen, Hilden, Germany). Collected resin was washed with 10 mL of 25 mM imidazole in PBS followed by 40 mM imidazole in PBS, and protein was eluted by incubation with 500 µL 300 mM imidazole in PBS twice. Protein was concentrated in a 30 kDa cut-off Amicon centrifugal concentrator (Millipore) to a final volume of ~50 µL. Final protein concentration was determined by BCA assay.

The coding sequence of human ST6GALNAC2 (58-374) was cloned into a pGEn2-DEST vector (DNASU, Arizona, US) containing an N-terminal triple His-Avi-sfGFP tag for purification and detection followed by a TEV cleavage site. The construct is preceded by a

TCM signal peptide to direct protein secretion into the media. For expression, 20 µg of endotoxin-free maxiprep DNA (ZymoPURE™ II Plasmid Maxiprep Kit, Zymo Research, US) was transfected into 20 mL of Expi293F® cells (Gibco®) using ExpiFectamine™ 293 Transfection kit (Thermo Fisher) and following the manufacturer's protocol. After 4 days incubation at 37°C, the media was collected and cleared by centrifugation. The recombinant protein was purified via its GFP-tag by incubating the media with 100 µL of home-made GFP-Clamp Agarose beads[8] for 2 hours at 4°C while mixing. Unbound fractions were collected by centrifugation, beads washed with 1 mL Buffer A (20 mM Hepes pH 7.5; 300 mM NaCl) and protein eluted from the beads by incubating for 4 hours in 100 µL Buffer A supplemented with 20 µL 1 mg/mL TEV protease (Kapust et al).[9] Elution fractions were collected by centrifugation and beads washed twice with 100 µL Buffer A. Elution steps were repeated twice. Fractions containing the cleaved protein were pooled, diluted 20-fold in Freezing Buffer (10mM Hepes, pH7; 50mM NaCl; 10% Glycerol) and concentrated to 300 µL using a Vivaspin®6 10K centrifugal tube (Sartorius, Göttingen, Germany). A concentration of 0.169 mg/mL was determined by NanoDrop sample was flash frozen and stored at -80°C until further use.

The human TMPRSS2 ectodomain (Uniprot identifier O15393, residues 111-492) was synthesised and cloned by Genscript into pcDNA 3.1(+) with a *Gaussia* luciferase secretion signal followed by a FLAG tag added on the N-terminus and the strep-tag on the C-terminus. It was expressed exactly as described above for the spike, except it was harvested on the sixth day post-transfection and purified from the supernatant with Streptactin XT (IBA Lifesciences GmbH, Germany).

##### *In vitro* glycosylation, CuAAC reaction and Western Blotting of 2P-WT, P681H, and P681R SARS-CoV-2 spike protein by WT or BH GalNAc-T1 & T2

UDP-GalN6yne was a gift from Junwon Choi and Carolyn Bertozzi.[1] *In vitro* glycosylation reactions of recombinant spike were performed in PCR tubes using 1 µg spike (2P-WT, P681H or P681R) in 20 µL final reaction volume containing 20.8 mM Tris-HCl pH 7.5, 50 mM NaCl, 10 mM $MnCl_2$ ($MnCl_2{\bullet}4H_2O$ ; Sigma Aldrich), 50 µM UDP-GalN6yne, 100 µM Uridine 5′-diphospho-N-acetylgalactosamine disodium salt (UDP-GalNAc; Sigma Aldrich) and 0.5 U Invitrogen® calf intestinal alkaline phosphatase (CIAP; Thermo Fisher Scientific) at 37 °C for 16 h using soluble WT or BH-GalNAc-T1/T2 at a final concentration of 100 nM. After incubation, reactions were heat inactivated at 95 °C for 20 s, then rapidly chilled to 4 °C. Crude reaction mixtures were subjected to CuAAC reactions in 30 µL final reaction volume containing 1.2 mM BTTAA (Jena Bioscience; Germany), 0.6 mM $CuSO_4$ ($CuSO_4{\bullet}5H_2O$ Sigma Aldrich), 10 mM L-Ascorbic acid sodium salt (Acros Organics; Geel, Belgium), 10 mM aminoguanidinium chloride (Cayman Chemical Company; Ann Arbor, USA), and 100 µM biotin picolyl azide (Sigma-Aldrich) for 18 h at room temperature with a 60 nM final spike concentration.

A 20 µL aliquot of each CuAAC sample (~600 µg Spike) was mixed with 6 µL 4X Protein Sample Loading Buffer (LI-COR Bioscience; *important*: *no reducing agents added since this abrogates the nanobody's epitope in the RBD*), heated at 90 °C for 3 min and samples subjected to SDS-PAGE on a 4–20% 18-well Criterion™ TGX™ Precast Midi Protein Gel (Bio-Rad Laboratories, UK) in Tris/Glycine/SDS buffer (Bio-Rad Laboratories) at 160 V for 80 min. Gels were transferred to a Trans-Blot Turbo RTA Midi 0.2 µm Nitrocellulose membrane (Bio-Rad Laboratories) using the Trans-Blot Turbo Transfer System according to the manufacturer's instructions and then left to dry to maximize protein retention. Membranes were rehydrated in MilliQ water followed by incubation in 5 mL Revert™ 700 Total Protein Stain (LI-COR Bioscience) for 5 min. Membranes were then washed twice with 5 mL Revert™ 700 wash and visualised on an Odyssey CLx® Imaging System (LI-COR Bioscience; Lincoln, USA). To identify biotinylation, membranes were first incubated in protein-free TBS blocking buffer (LI-COR Bioscience) for 1 h and then incubated in IRDye® 800CW Streptavidin (LI-COR Bioscience) according to the manufacturer's instructions.

Membranes were washed four times with TBST, once with TBS for 5 min each, and then scanned by CLx® Odyssey. Then, membranes were counterstained with 10 mL of the mouse-Fc nanobody (MsFc-NbA5, 1:5,000 in blocking buffer) produced in-house at 4 °C overnight (alternatively, for 1 h at room temperature). Membranes were washed three times with 1X Tris-Buffered Saline, 0.1% Tween® 20 Detergent (TBST) and once with TBS, for 5 min each. Detection was performed with an IRDye 680RD goat anti-mouse IgG solution in protein-free TBS blocking buffer (LI-COR Bioscience) for 30 min at room temperature. Membranes were washed three times with TBST and once with TBS as before and then scanned by CLx® Odyssey.

##### Azido ITag ($N_3$-ITag) CuAAC and engineered cell sample processing for mass spectrometry

25 µL aliquots containing 2 µg recombinant S from BH-T1 or BH-T2 engineered cells (see above) in 20 mM Tris pH 8.0, 150 mM NaCl buffer were treated with PNGase-F (Promega, 5 µL of a 1:10 dilution in PBS) for 3 h at 37 °C. CuAAC reactions were performed on the PNGase-F treated mixtures in a 45 µL final reaction volume containing 2.4 mM BTTAA (Jena Bioscience; Germany), 1.2 mM $CuSO_4$ ($CuSO_4$•$5H_2O$ Sigma Aldrich), 10 mM L-Ascorbic acid sodium salt (Acros Organics; Geel, Belgium), 10 mM aminoguanidinium chloride (Cayman Chemical Company; Ann Arbor, USA), and 1 mM $N_3$-ITag for 16 h at room temperature. Reaction mixtures were then dried in a centrifugal concentrator and redissolved in 20 µL DI $H_2O$. Each CuAAC sample was mixed with 6 µL 4X Protein Sample Loading Buffer (LI-COR Bioscience; *important*: *no DTT or other reducing agents added since this abrogates the nanobody's epitope in the RBD*), heated at 90 °C for 3 min and samples subjected to SDS-PAGE on a 4–20% 18-well Criterion™ TGX™ Precast Midi Protein Gel (Bio-Rad Laboratories, UK) in Tris/Glycine/SDS buffer (Bio-Rad Laboratories) at 160 V for 80 mins. Gels were stained with SafeBlue Protein Stain (NBS Biologicals Huntingdon, UK) for 30 min. Protein bands corresponding to full length S and combined S1/S2 were excised from the gel with the help of a clean scalpel, placed into a microfuge tube, cut into ca. 1x1 mm pieces, and spun down in a benchtop microcentrifuge. Gel pieces were destained with 100 µL of 100 mM Seppro™ Ammonium Bicarbonate Buffer (Ambic; Merck) 1:1, vol/vol in Pierce™ Acetonitrile LC-MS Grade (MeCN; Thermo Fisher Scientific) and incubated for 30 min with occasional vortexing. This step was repeated three more times for 10 min each, discarding the supernatant after every wash. Then, 400 µL of MeCN were added to the gel pieces and incubated at room temperature with occasional vortexing until pieces had shrunk and were white (ca. 10 min). Gel pieces were subjected to reduction with 10 mM DL-Dithiothreitol (Sigma Aldrich) in 100 mM Ambic and incubated at 56 °C for 45 min. After cooling to room temperature, all liquid was removed followed by addition of 55 mM iodoacetamide (Sigma Aldrich) in 100 mM Ambic buffer and incubated for 30 min in the dark at room temperature. The supernatant was removed and washed three times with 100 mM Ambic buffer. The gel pieces were incubated with MeCN for 10 min, the supernatant was discarded, and samples were dried in a centrifugal concentrator.

For proteolytic digestion, gel pieces were rehydrated with a 25 ng/µL Glu-C (Sequencing Grade; Promega, UK) solution in 100 mM Ambic and left at 37 °C to saturate for a few min. Once rehydrated, added enough 100 mM Ambic to cover the gel pieces and then left to digest for 2 h at 37 °C. After 2 hour, a volume equal to that of the 25ng/µL Glu-C solution, of a 25 ng/µL Trypsin Gold (Mass Spectrometry Grade; Promega, UK) solution in 100 mM Ambic was added to the samples and left incubating at 37 °C for 16 h. Samples were then centrifuged, the supernatant ((glyco-)peptide fraction) was collected and transferred to a low-bind tube. To further extract the peptides, 20 µL of extraction buffer (*i.e.* 1:1 (vol/vol) 2% formic acid (FA)/MeCN) were added to each tube with the gel pieces and incubated for 15 min at 37 °C in a shaker. Supernatants were collected and added to the low-bind tube containing the first portion of the digest. Digests were dried in a centrifugal vacuum concentrator (can be safely stored at -20 °C for a couple of months).

#### Proteomics analysis

Dried (glyco-)peptide fractions were resuspended in 15 µL of 0.1% (v/v) FA in LCMS-grade water, sonicated for 15 min, vortexed briefly and centrifuged for 5 min at 18,000 g. Sample mixtures were analyzed by nanoflow LC-MS/MS using an Orbitrap Eclipse with ETD (Thermo Fisher) coupled to an UltiMate 3000 RSLCnano (Thermo Fisher). The sample (10 µL out of 15 µL for peptide fractions) was loaded via autosampler isocratically onto a 50 cm, 75 µm PepMap RSLC C18 column (ES903) after pre-concentation onto a 2 cm, 75 µm Acclaim PepMap100 m nanoViper. The column was held at 40 °C using a column heater in the EASY-Spray ionization source (Thermo Fisher). The samples were eluted at a constant flow rate of 0.275 µL/min using a 120- and 140-min gradient for peptides and glycopeptides, respectively. Solvents were 5% (v/v) DMSO, 95% (v/v) 0.1% FA in water (A) and 5% (v/v) DMSO, 20% (v/v) 0.1% FA in water, 75% (v/v) 0.1% FA in MeCN (B).

The gradient profile was as follows: 0 min: 2% B; 6 min: 2% B; 114 min: 40% B; 115 min: 5% B; 119 min: 95% B; 120 min: 2% B; 140 min: 2% B.

For glycopeptide identification, MS1 scans were collected with a mass range from 300-1500 m/z, 120K resolution, 4e5 ion inject target, and 50 ms maximum inject time. Dynamic exclusion 21 was set to exclude for 10 s with a repeat count of 3. Charge states 2-6 with an intensity greater than 1e4 were selected for fragmentation at top speed for 3 s. Selected precursors were fragmented using HCD at 28% nCE with 2 Da isolation window, 5e4 inject target, and 54 ms maximum inject time before collection at 30K resolution in the Orbitrap. For precursors from 300-1000 m/z, presence of 3 oxonium ions over 5% relative abundance triggered a charge calibrated ETD scan to be collected in the ion trap with a 3 Da isolation window, 1e4 inject target, and 100ms maximum injection time.

Data evaluation of glycopeptides was performed with Byonic™ (Protein Metrics, Cupertino, USA). For glycopeptide analysis, search parameters included semi-specific cleavage specificity at the C-terminal site of R and K, with two missed cleavages allowed. Mass tolerance was set at 10 ppm for MS1s, 20 ppm for HCD MS2s, and 0.2 Da for ETD MS2s. Carbamidomethyl cysteine was set as a fixed modification. Variable modifications included methionine oxidation (common 1), asparagine deamidation (common 1), and a custom database of O-glycans that included HexNAc, HexNAc-NeuAc, HexNAc-Hex, HexNAc-Hex-NeuAc, $HexNAc_2$-Hex-NeuAc and HexNAc-Hex-$NeuAc_2$ with an additional 287.1371 m/z to account for the chemical modification. A maximum of two variable modifications were allowed per peptide. All identifications with |logP| greater than 3 that contained chemically modified glycans were manually validated and localized using a combination of HCD and ETD information.

#### Solid phase synthesis of peptides

Solid phase synthesis of peptides took place on an automated peptide synthesizer (Activotec, P11) using a Rink Amide AM resin LL resin (0.05 mmol; Merck), Fmoc-Ser(tBU)-wang-LL resin (0.05 mmol; Merck) and N(α)-Fmoc amino acids, including Fmoc-Tyr(3-$NO_2$)-COOH,Fmoc-Abz-COOH or Fmoc-Thr(GalNAc(Ac)$_3$)-OH as appropriate. HATU was used as the coupling reagent with 5- fold excess of amino acids.

Peptides were cleaved from the resin and protecting groups removed by addition of a cleavage solution (95% (v/v) trifluoroacetic acid (TFA), 2.5% (v/v) water, 2.5% (v/v) triisopropylsilane, TIS). If peptides contained cysteine or methionine, ethane-1,2-dithiol (EDT) was also added. (92.5% (v/v) TFA, 2.5% (v/v) water, 2.5% (v/v) TIS, 2.5% (v/v) EDT). Those peptides containing multiple arginines were cleaved using bromotrimethylsilane (TMSBr) or Phenol. After 2 h, the resin was removed by filtration and peptides were precipitated with diethyl ether on ice. Peptides were isolated by centrifugation, then dissolved in water and freeze dried overnight. Portions of the peptides were purified on a C8 reverse phase HPLC column (Agilent PrepHT Zorbax 300SB-C8, 21.2x250 mm, 7 m) using a linear solvent gradient of 10-50% MeCN (0.08% TFA) in water (0.08% TFA) over 40 min at a flow rate of 8 mL/min. The peptide containing Thr(GalNAc(Ac)$_3$) was deacetylated with 5% hydrazine monohydrate in water for 1h and directly purified as above. The purified peptides were analysed by LC–MS on an Agilent 1100 LC-MSD. The calculated molecular weights of the peptides were in agreement with the masses found.

Peptides used were $NH_2$-ASYQTQTNSPRRARS-$CONH_2$ (WT), $NH_2$-ASYHTQTNSPRRARS-$CONH_2$ (Q675H), $NH_2$-ASYRTQTNSPRRARS-$CONH_2$ (Q675R), $NH_2$-ASYHTHTNSPRRARS-$CONH_2$ (Q675H Q677H) $NH_2$-ASYHTRTNSPRRARS-$CONH_2$ (Q675H Q677R), $NH_2$-ASYQTHTNSPRRARS-$CONH_2$ (Q677H), $NH_2$-ASYQTRTNSPRRARS-$CONH_2$ (Q677R), $NH_2$-ASYQTQTKSPRRARS-$CONH_2$ (N679K), $NH_2$-ASYQTQTNSHRRARS-$CONH_2$ (P681H), $NH_2$-ASYQTQTNSRRRARS-$CONH_2$ (P681R), Abz-ASYQTQTNSPRRARSVAS-Tyr(3-$NO_2$)-$CONH_2$ (**FRET-1**), Abz-ASYQTQTNSHRRARSVAS-Tyr(3-NO2)-$CONH_2$ (**FRET-2**), Abz-ASYQTQT(GalNAc)NSPRRARSVAS-Tyr(3-$NO_2$)-$CONH_2$ (**FRET-3**), Abz-ASYQTQTNSNSPRRAR-COOH (**SI-1**). High resolution mass spectrometry (HRMS) was taken with a Synapt G2-Si (Waters, Milford, USA). HPLC samples were run on Acquity qDA UPLC-MS (Waters), Diode Array spectrum 200-400 nm, ACQUITY UPLC® BEH C18 column (1.7 µm, 2.1 x 50 mm) and flow rate 0.5 mL/min using Buffer A: water containing 0.1% formic acid and Buffer B: MeCN containing 0.1% formic acid, linear gradient from 3% B to 97% buffer B over 4 min.

#### Glycosylation of a SARS-CoV-2 peptide panel

Glycosylation reactions were performed in 50 µL volume containing Tris HCl pH 7.5 (20 mM), NaCl (50 mM), $MnCl_2$ (10 mM), UDP-GalNAc (250 µM), peptide (100 µM) and recombinant WT-GalNAc-T1 (790 nM). Reaction mixtures were gently pipetted to assure a homogeneous distribution of all the components and was placed in a static incubator at 37 °C for 18 h. Reactions were stopped by addition of MeCN (20 µL) and freezing at -20 °C for 2 h. The reactions were centrifuged to remove any precipitated enzyme pellet and the supernatant was loaded into a 60 mg Strata-X cartridge (Phenomenex, Torrance, USA) pre-conditioned sequentially with MeCN and water containing 0.1% (v/v) FA. The column was washed with water containing 0.1% (v/v) FA, and peptide/glycopeptide mixtures were loaded and eluted with 80% (v/v) aqueous MeCN with 0.1% (v/v) FA. Eluted samples were freeze-dried to remove the solvent and resuspended in 50 µL of water. A 10 µL aliquot was diluted 1:1 with water containing 0.1% (v/v) FA and characterized by Ultra Performance Liquid Chromatography Mass Spectrometry (UPLC-MS) analysis using a Waters Acquity Ultra Performance System (Waters) equipped with ACQUITY UPLC® BEH C18 column (1.7 µm, 2.1 x 50 mm) and flow rate 0.5 mL/min using Buffer A: water containing 0.1% (v/v) FA) and Buffer B: MeCN containing 0.1% FA. Samples were run on a linear gradient from 3% B to 97%B over 4 min.

Data were recorded from three independent replicates, by extraction of total ion count of the peak corresponding to $[M]^{+3}$ of starting material, glycosylated and di-glycosylated product and their integration as established previously.[10]

#### Synthesis of FRET-4 through glycosylation of FRET-1 with GalN6yne

Glycosylation was performed in 1.085 mL Tris HCl pH 7.5 (20 mM), NaCl (50 mM), $MnCl_2$ (10 mM) containing UDP-GalN6yne (300 µM), peptide **FRET-1** (200 µM, 500 µg) and recombinant BH-GalNAc-T1 (292 nM). The reaction mixture was incubated at 37 °C for 18 h. Completion of the reaction was monitored by UPLC-MS. The reaction was stopped by addition of MeCN (200 µL) and kept at -20°C for 2 h. After thowing the reaction mixture was centrifuged to remove any precipitate and the supernatant was desalted using Strata-X cartridge. The eluted fraction was freeze dried. After resuspension in minimal volume of water with 0.1% (v/v) FA and MeCN 1:1 (v/v), the glycopeptide was purified by HPLC on a 1260 Infinity II MDAP system (Agilent Technologies, UK) equipped with an XBridge® BEH Amide OBD™ Prep column (130Å, 5 µm, 10 mm x 100 mm) and the following solvent system: Buffer A:10 mM Ammonium formate pH 4.5; Buffer B: 10 mM Ammonium formate in MeCN: water 90:10.The glycopeptide was eluted with a linear gradient from 90% Buffer B to 55% Buffer B over 15 min at a flow rate of 19 mL/min and collected by detection of UV absorbance at 210 nm and/or positive mode mass signal corresponding expected molecular mass of 2560.2055 Fractions were freeze dried and dissolved in water (200 µL). Isolated glycopeptide was quantified by nanodrop ($\lambda_{abs}$ 280 nm) against a standard curve of peptide **FRET-4** and determined to be 185 nmol (85% yield). The purity and structure of the product was confirmed by UPLC-MS and HRMS. HRMS (m/z): $[M+3H]^{3+}$ calcd.for C108H165N35O38, 854. 4018; found: 854.4103.

##### Synthesis of FRET-5 and FRET-6 through CuAAC-facilitated functionalization of FRET-4

CuAAC reactions were performed in 30 µL total volume of 25 mM Tris Cl pH 8, 50 mM BTTAA, 60 mM $CuSO_4$ and 100 µM sodium ascorbate. To the mixture, 100 µL of a 200 µM glycopeptide **FRET-4** 200 µM were added, followed by 20 µL of a 10 mM stock solution of either 6-azido glucose or azido-propionic acid (Sigma Aldrich, UK) in water. The reactions were incubated at 37 °C for 3 h and checked by UPLC-MS to confirm quantitative conversion. Reaction mixtures were subjected to STRATA-X cartridge purification to yield the glycopeptides **FRET-5** and **FRET-6** in good purity. The eluted fractions were freeze dried, resuspended in 100 µL and quantified by nanodrop ($\lambda_{abs}$ 280 nm) against a standard curve of peptide **FRET-4** The purity and structure of the product was confirmed by UPLC and HRMS. MW of FRET-5: 2765.2753, FRET-6: 2675.2437 **HRMS** (m/z): or **FRET-5** $[M+3H]^{3+}$calcd. for C114H176N38O43, 922.7584, found: 922.7681; for **FRET-6** $[M+4H]^{4+}$ calcd. for C111H170N38O40, 669.8109, found: 669.8206.

##### Preparation of FRET-7 via chemoenzymatic galactosylation of FRET-3

*Dm*C1GalT1 was used for the *in vitro* galactosylation reaction of FRET-3.[11] Synthetic **FRET-3** peptide (500 µM, 3.8 mg), UDP-Gal (750 µM) and *Dm*C1GalT1 (1 µM) in Buffer containing Tris pH 7.5 (25 mM), NaCl (150 mM), $MnCl_2$ (0.05 mM) and BSA (bovine serum albumin) (1 mg/mL) were mixed in total volume of 3 mL and placed in a shaking incubator at 220 rpm and 37 °C for 16 hours. The reaction was monitored by UPLC-MS (Waters) equipped with ACQUITY UPLC® BEH C18 column (1.7 µm, 2.1 x 50 mm). A sample (3 µL) of the reaction mixture was run on a linear gradient from 3% B to 97% B over 4 min at 0.5 mL/min flow rate to assess the progress of the reaction. After 16 hours all starting material was consumed and two products observed- a mono-galactosylated product (67%) and a doubly-galactosylated peptide (33%). The reaction mixture was quenched by the addition of equivalent volume of cold MeCN (3 mL), left on ice for 30 min and centrifugated at 1300 rpm and 4 °C for 30 min to facilitate the precipitation of the enzyme. The supernatant was lyophilised, and the dried product desalted by passing it through a pre-conditioned 60 g STRATA-X cartridge and eluted with 80% MeCN containing 0.1 % formic acid. To achieve isolation of the target monogalactosylated peptide, the Strata-X fraction was further purified on Agilent 1260 Infinity II MDAP system (Agilent Technologies, UK) equipped with Agilent Prep- C18 column (100Å, 5 µm, 21.2 mm x 50 mm) and the following solvent system: Buffer A: water containing 0.1% formic acid; Buffer B: MeCN containing 0.1% formic acid. The target glycopeptide was eluted with gradient of 2-25% Buffer B over 10 min at a flow rate of 25 mL/min. Fractions containing the product were collected by automatic fraction collector set to trigger collection at 210 nm UV absorbance and/or positive mode mass signal, corresponding to the expected molecular mass of the 2670.2270 Da. The pure fractions were pooled and freeze dried to yield 1.9 mg of pure product (**FRET-7)**, 46%. The purity and structure of the product was confirmed by UPLC and **HRMS** (m/z) $[M+3H]^{3+}$ calcd for C110H171N35O43, 891.0756, found: 891.0823.

##### Synthesis of FRET-8 via chemoenzymatic sialylation of FRET-3

Synthetic **FRET-3** peptide (1 mM, 2.5 mg), CMP-Neu5Ac (1.5 mM) and ST6GALNAC1 (Glyco Expression Technologies, USA) (150 µg/mL) in Buffer containing Tris pH 7.5 (100 mM), $MgCl_2$ (10 mM) and CIAP (10 U/mL) were mixed in a total volume of 1 mL and placed in a shaking incubator at 220 rpm and 37°C for 16 hours. Completion of the reaction was monitored by taking a sample of the reaction mixture (3 µL) and running it on UPLC-MS (Waters). After 16 hours, 95% of the starting material was converted to the sialylated product. The reaction mixture was quenched, and the product purified in an identical manner to **FRET-7**. Fractions containing the product were collected by automatic fraction collector set to trigger collection at 210 nm UV absorbance and/or positive mode mass signal, corresponding to the expected molecular mass of the 2799.2-696 Da. The pure fractions were pooled and freeze-dried to yield 1.4 mg of pure product (**FRET-8)**, 50%. The purity and structure of the product was confirmed by UPLC and **HRMS** (m/z) $[M+4H]^{4+}$ calcd. for C115H178N36O46, 700.8174, found: 700.8279.

##### Synthesis of FRET-9 via chemoenzymatic sialylation of FRET-7

**FRET-7** peptide (20 µM, 160 µg), CMP-Neu5Ac (50 µM) and ST6GALNAC2 (10 µg/mL) in Buffer containing Tris pH 7.5 (100 mM), $MgCl_2$ (10 mM) and CIAP (10 U/mL) were mixed in a total volume of 3 mL and placed in a shaking incubator at 220 rpm and 37°C for 48 hours. Reaction was monitored by UPLC-MS. After 48 hours, 89% of the starting material was converted to the sialylated product. The reaction mixture was quenched, and the product purified in an identical manner to FRET-7 and FRET-8. Fractions with mass peak corresponding to the expected molecular mass of the 2961.3224 Da were pooled and freeze-dried to yield 0.4 mg of pure product (**FRET-9)**, 30%. The purity and structure of the product was confirmed by UPLC and **HRMS** (m/z) $[M+4H]^{4+}$ calcd. for C121H188N36O51, 741.3306, mass found: 741.3380.

##### Furin cleavage FRET analysis with glycopeptides FRET-1 to FRET-6

Peptides used for the FRET analysis were dissolved in water and their concentration adjusted using calibration curve of the FRET-1 peptide based on the integration of the UV absorbance signal at 320 nm, measured by UPLC.

Reactions were set up on ice, in black frame/white wells 384-well plate (VWR, UK) in a final volume of 25 µL. Each well contained 5 µL of 100 µM peptide (FRET-1 to FRET -6) and either 20 µL of 1 U/mL human furin (New England Biolabs, Ipswitch, USA) in reaction buffer (100 mM HEPES pH 7.5, 0.5% (v/v) Triton-X 100, 1 mM, $CaCl_2$, 1 mM 2-mercaptoethanol) or 20 µL of buffer only as a negative control. The measurements were performed with an EnSight Multimode Plate Reader (PerkinElmer, UK) set at 30°C and with excitation wavelength $\lambda_{ext}$ of 320 and emission wavelength $\lambda_{ems}$ of 430 nm, taking one reading per minute for 90 minutes. Each of the independent experiments consisted of two technical replicates to compensate for the occasional abnormality caused by bubble formation during the set up. Each reaction was reproduced as at least three independent experiments.

The rate of cleavage was calculated by measuring the increase of fluorescence during the linear phase of the reaction (20-90 min). Each signal was normalised by subtracting the signal of samples lacking furin for each time point and setting the starting point to 0. Linear regression analysis was performed, and the slopes converted to rate of reaction using a calibration curve of a control peptide containing only the N-terminal fluorophore (**SI-1**). The calibration curve was plotted by measuring the fluorescence signal of a series of concentrations (0.625 µM to 20 µM) of the control peptide in the reaction buffer. The signal for each concentration was averaged between the 90 mins reaction time and three independent replicates. Graphs were plotted and statistical analysis performed using GraphPad Prism 9 (Dotmatics).

##### Furin cleavage FRET assay (FRET-1, FRET-3 and FRET-7 to FRET-9)

Reactions were set up on ice, in black frame/white wells 384-well plate (VWR, UK) in a final volume of 25 µL. Each well contained 10 µL of 14 µM peptide (FRET-1 to FRET -3 and FRET-7 to FRET-9) and either 15 µL of 2.7 U/mL human furin (New England Biolabs, Ipswitch, USA) in reaction buffer (100 mM HEPES pH 7.5, 0.5% (v/v) Triton-X 100, 1 mM, $CaCl_2$, 1 mM 2-mercaptoethanol) or 15 µL of buffer only as a negative control. The measurements were taken and data processed as above. The rate of cleavage was calculated by measuring the increase of fluorescence during the linear phase of the reaction (20-60 min). Each of the three independent experiments consisted of two technical replicates to compensate for the occasional abnormality caused by bubble formation during the set up. Where possible an average of both technical replicates was taken.

##### TMPRSS2 cleavage FRET assay (FRET-1 to FRET-3 and FRET-7 to FRET-9)

Reaction set up and measurements were performed as described above (furin assay) with exception to enzyme concentration. TMPRSS2 was at concentration of 7.4 µg/ml and the reaction buffer consisted of 0.1 M Tris-Cl pH 8, 0.15 M NaCl, 1 mM EDTA and 50 mM biotin. The linear phase of the reaction was 20-40 min. Each reaction was reproduced as three independent experiments.

##### Evaluation of T cell responses using the Interferon-gamma ELISpot assay

To ascertain the immunogenicity of this area of interest, we employed the Interferon-gamma ELISpot assay, that allows to detect T cell recognition of synthetic peptides in peripheral blood mononuclear cells (PBMC).

Frozen PBMC were obtained from the Crick-Legacy study, which is a SARS-CoV-2 longitudinal study designed to investigate the immune response to, and protection offered by Covid-19 vaccines in two prospective cohorts (patient-facing healthcare workers at UCLH and staff recruited from the Francis Crick Institute), established in January 2021 (NCT04750356), as described (Wall EC et al, Lancet. 2021, 19;397(10292):2331-2333).[12]

PBMC were thawed and seeded at 300,000/well using the Interferon-gamma ELISpot assay, using pre-coated plates from the Human IFN-γ ELISpotPRO kit (Mabtech 3420-2APT-10) as described (Fendler A et al, Nat Cancer. 2021, 2(12):1305-1320).[13]

Cells were stimulated overnight using SARS-CoV-2 peptide pools covering the S, M, N proteins (PepTivator peptide pools, Miltenyi Biotec, Surrey, UK) at 1 µg/ml per peptide, as well as the WT-GalNAc, P681H and WT (glyco)peptides (at 10µg/ml each). Results are expressed as IFN-gamma spot forming cells (SFC) per million PBMC following correction for the unstimulated background by subtraction.

### CHARACTERIZATION OF FRET (GLYCO-)PEPTIDE REPORTERS

HPLC traces and annotated HRMS spectra for synthetic peptides **FRET-1** to **FRET-9** and their corresponding multiply charged ions.

**FRET-1**:

HPLC

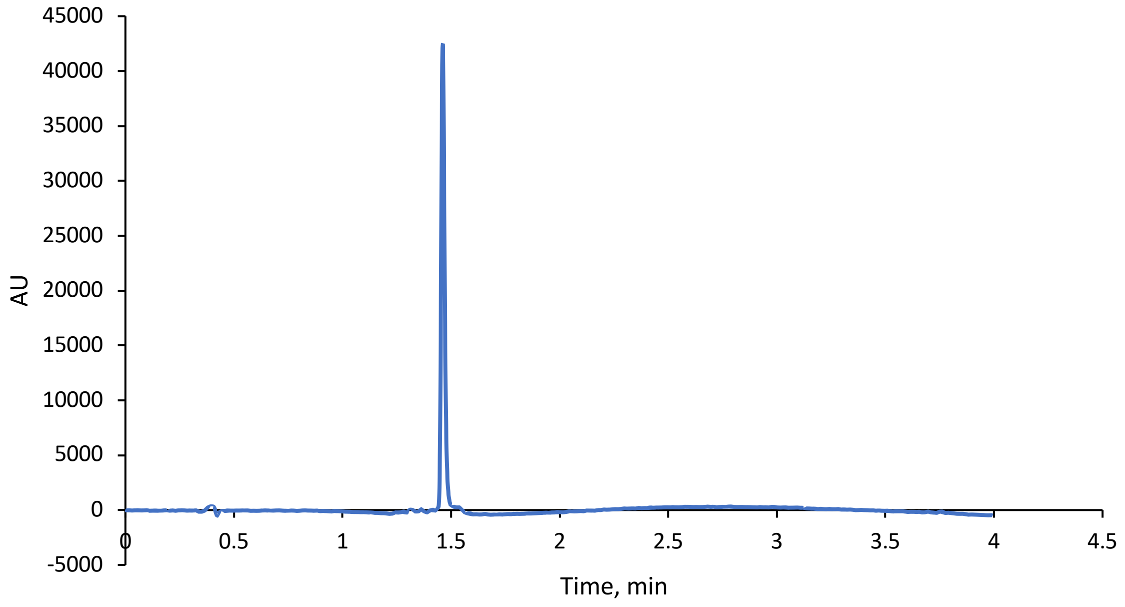

HRMS

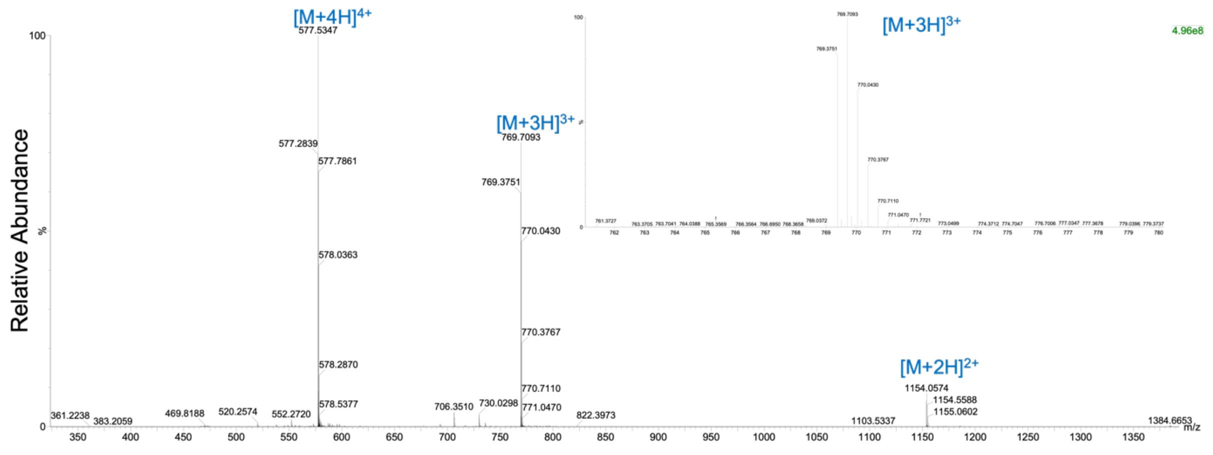

**FRET-2**:

HPLC

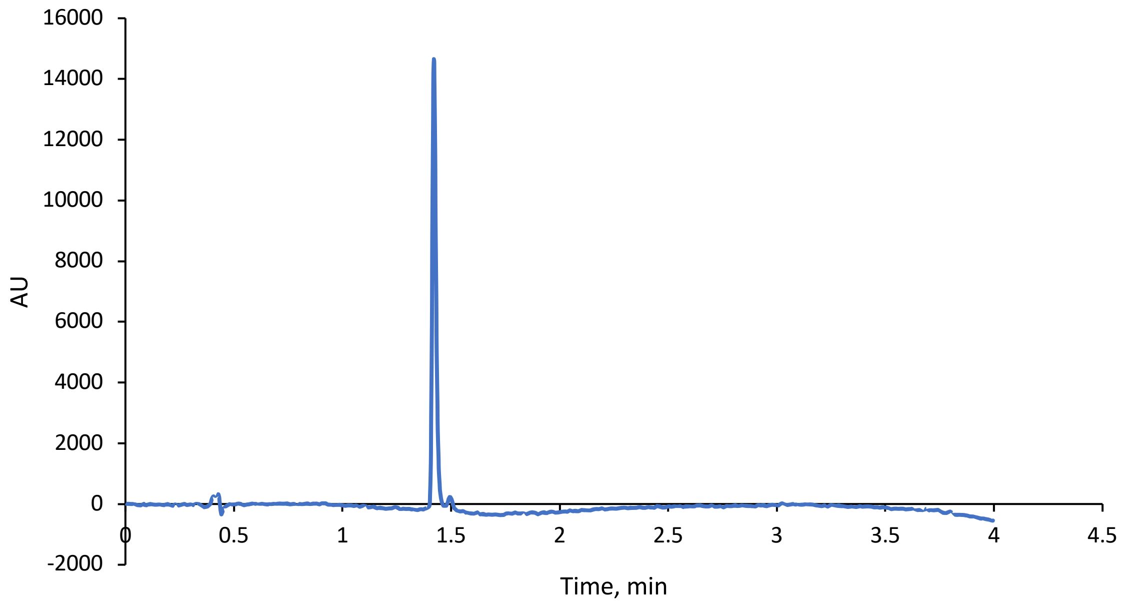

HRMS

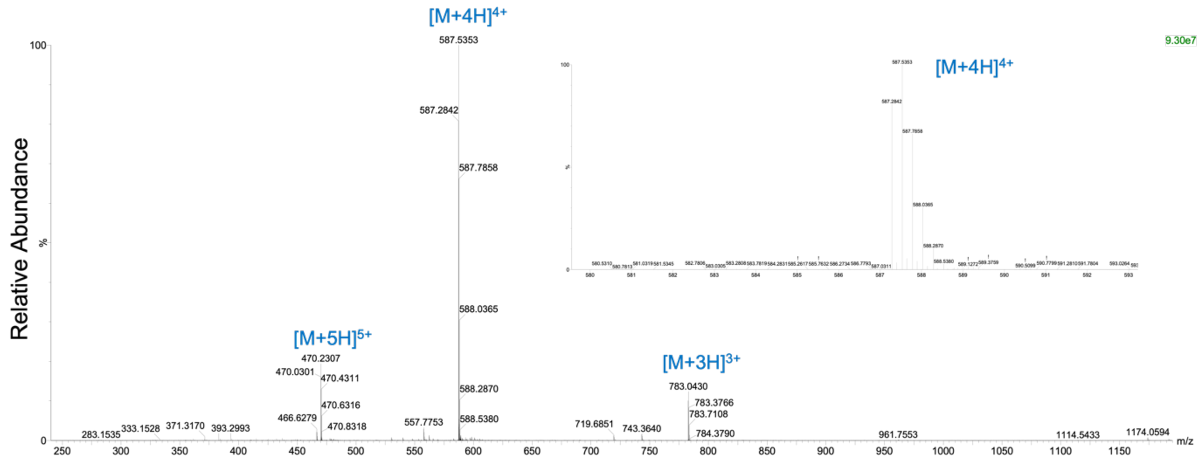

**FRET-3**:

HPLC

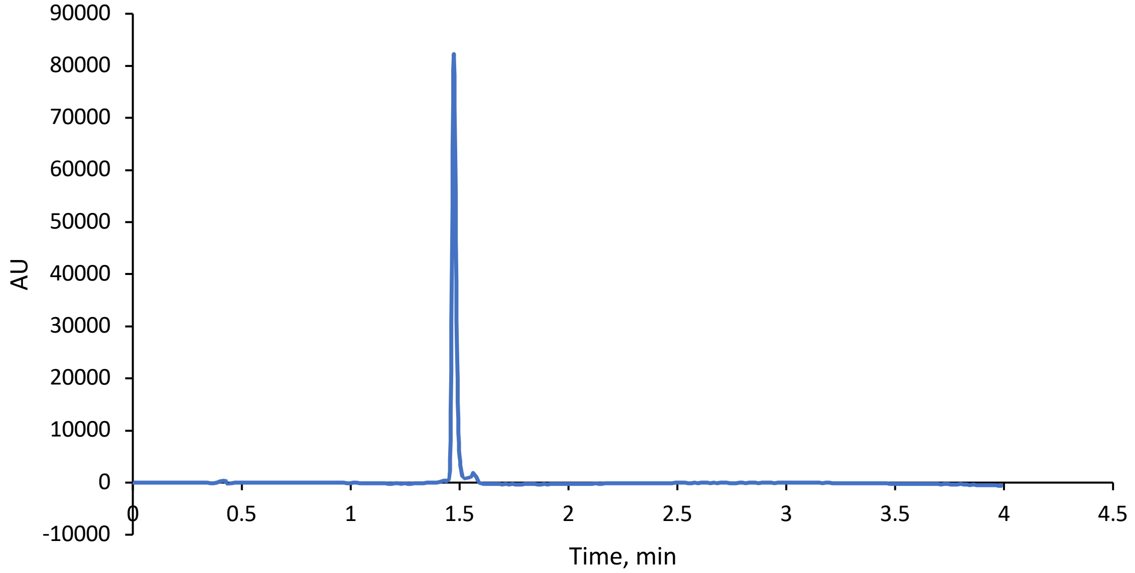

HRMS

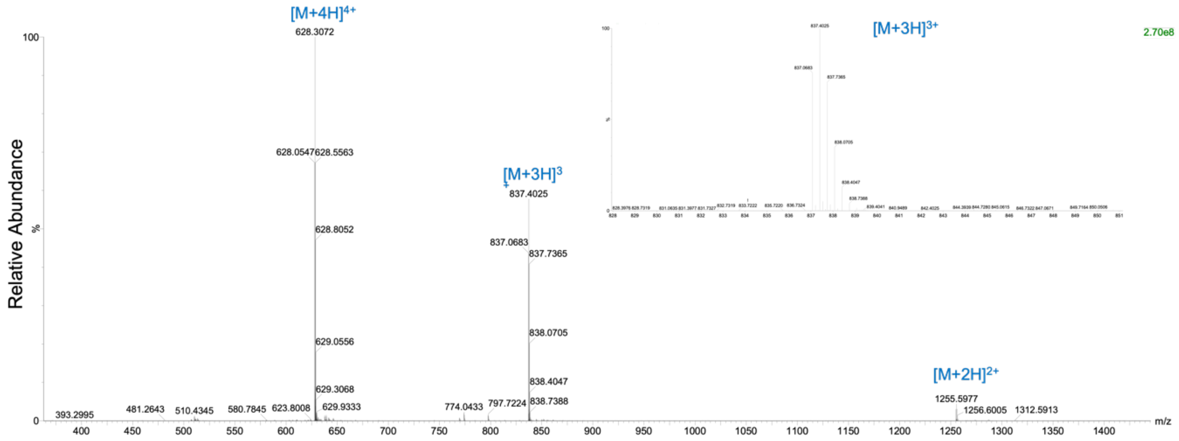

**FRET-4**:

HPLC

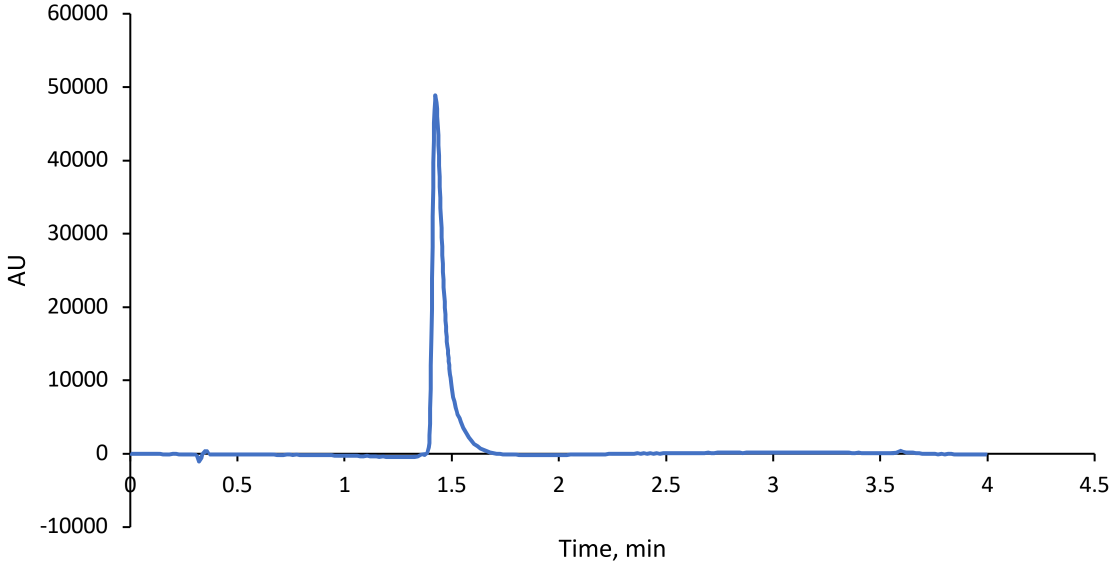

HRMS

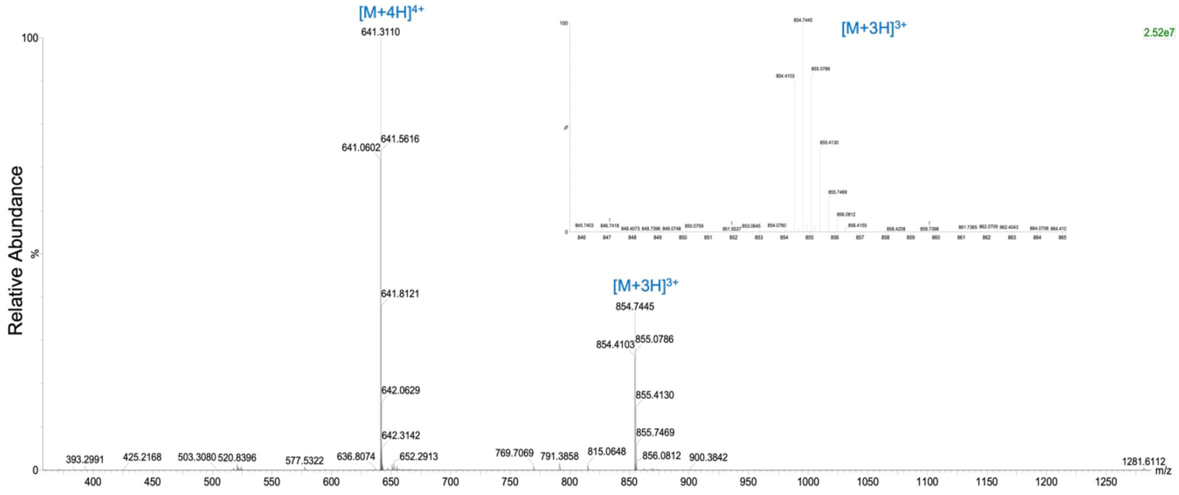

**FRET-5**:

HPLC

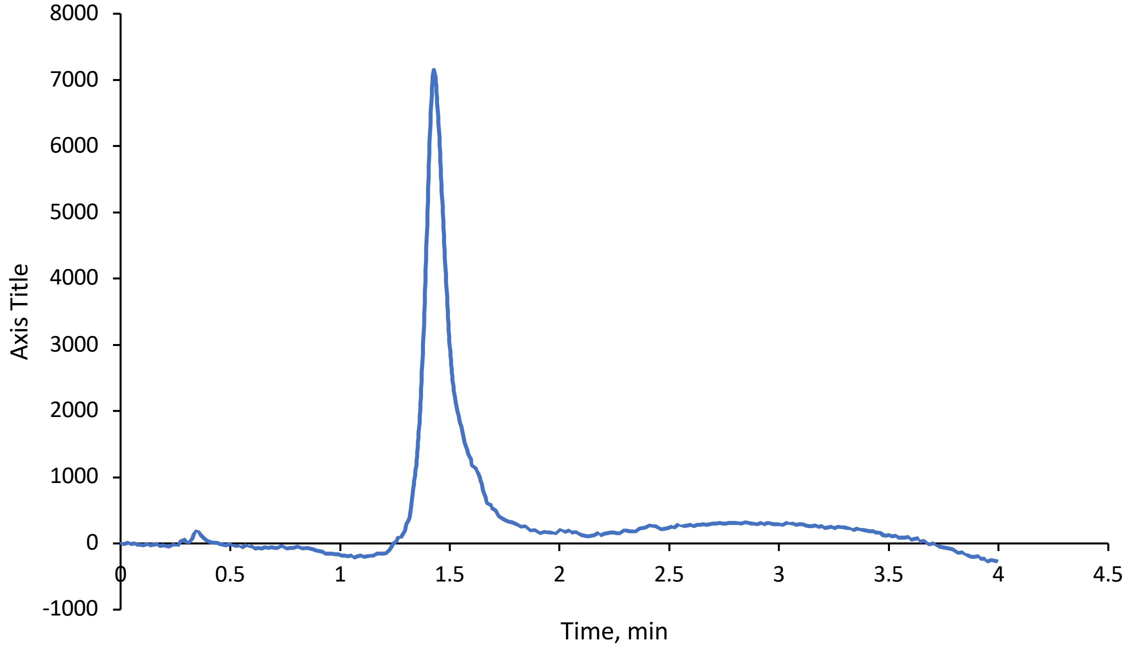

HRMS

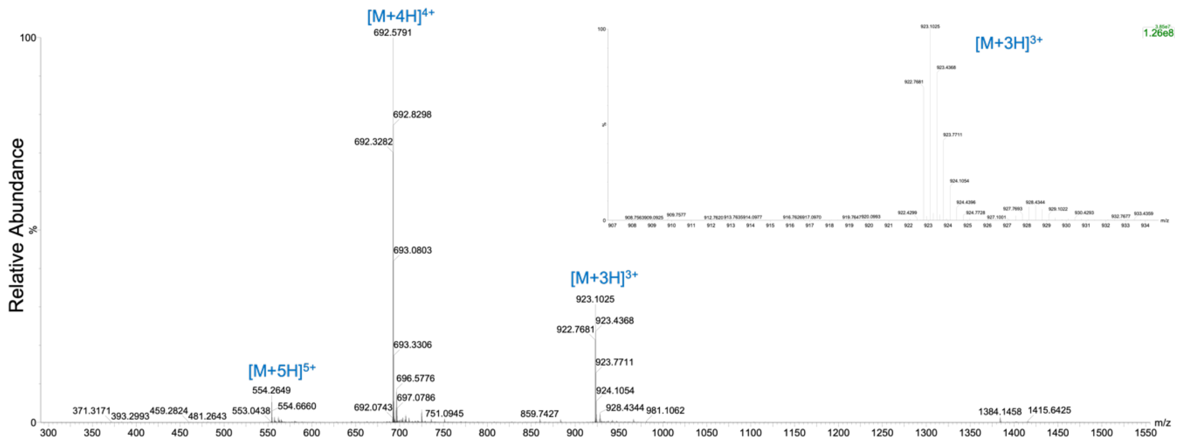

**FRET-6**:

HPLC

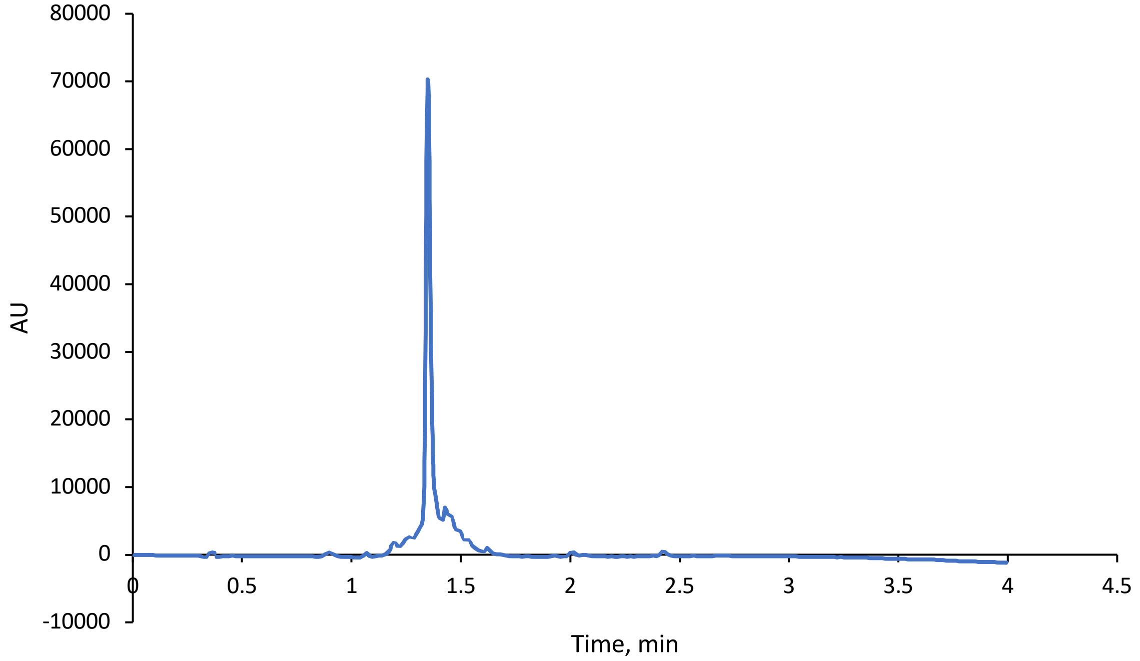

HRMS

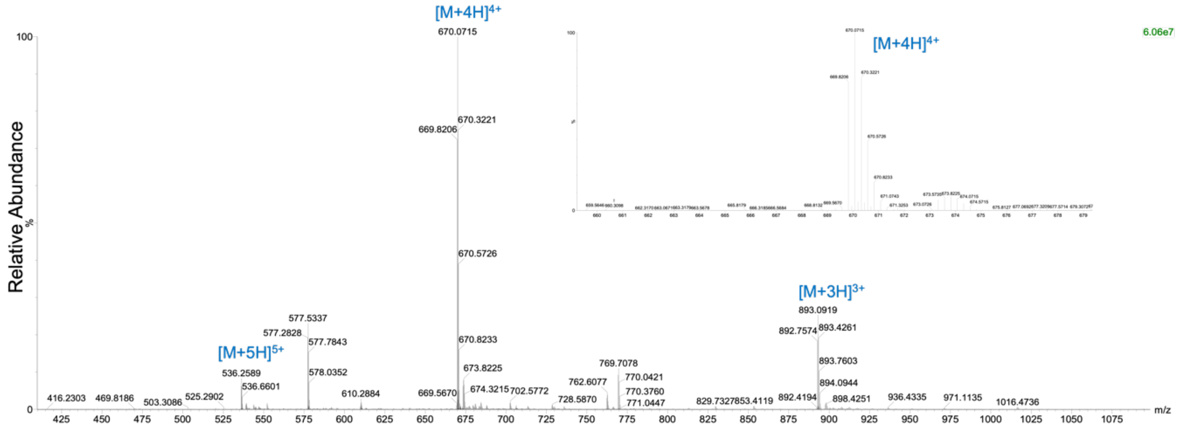

**FRET-7**:

HPLC

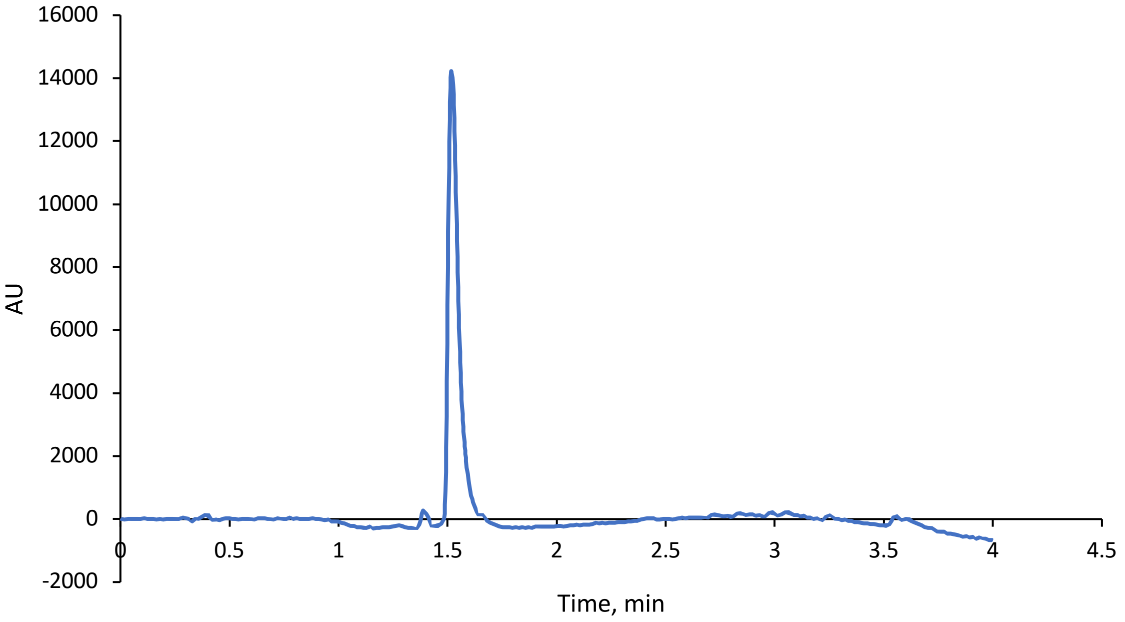

HRMS

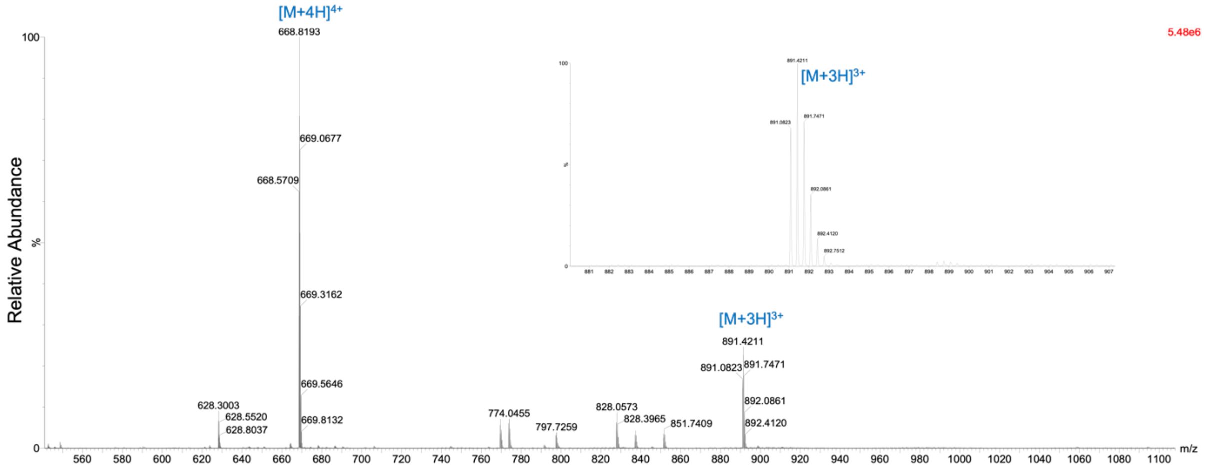

**FRET-8**:

HPLC

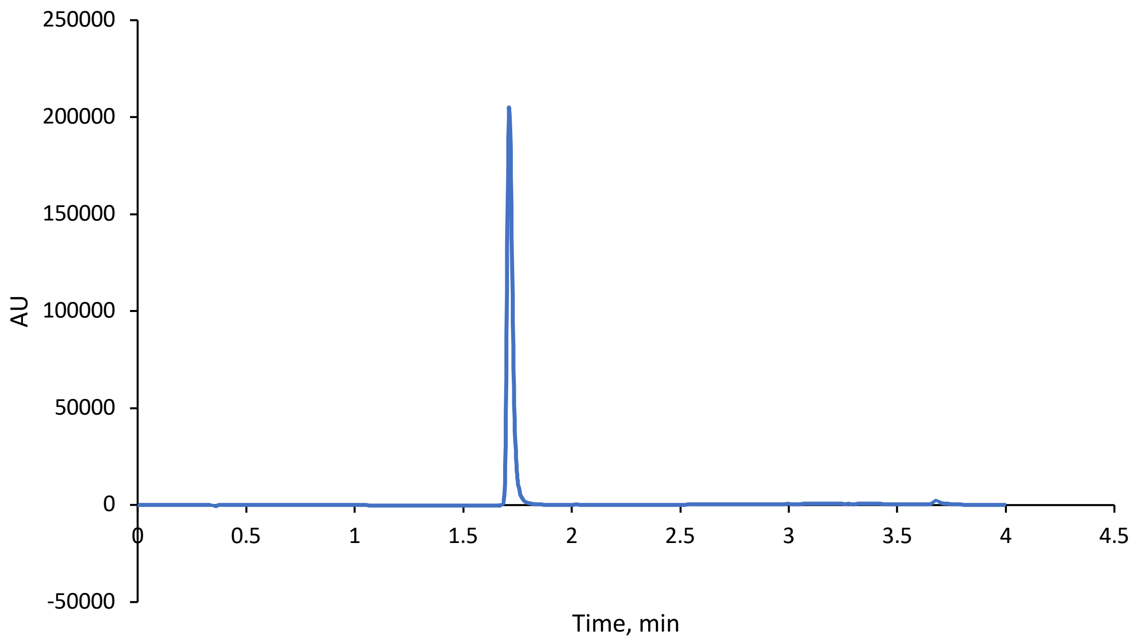

HRMS

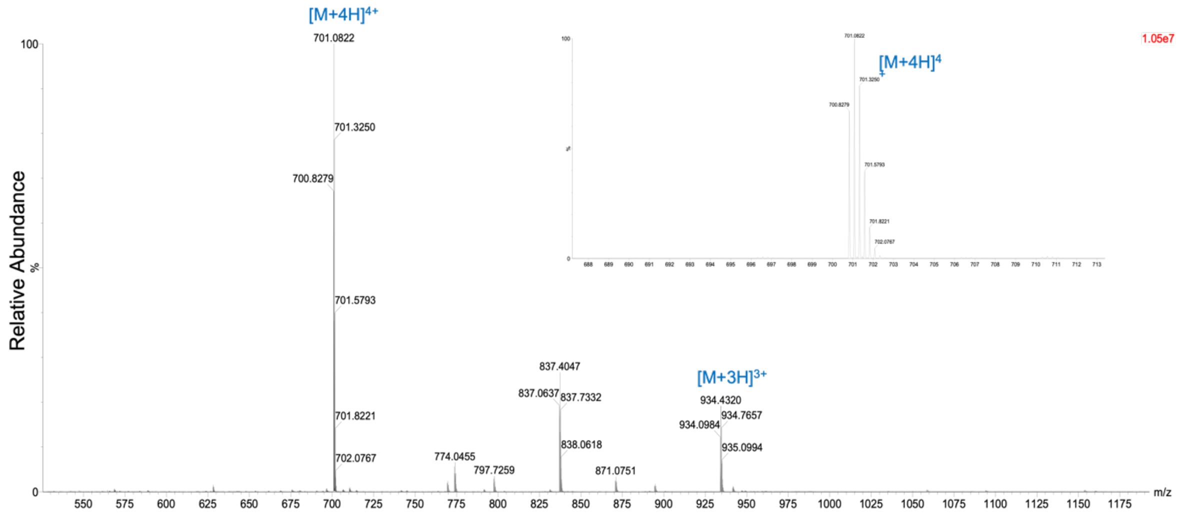

**FRET-9**:

HPLC

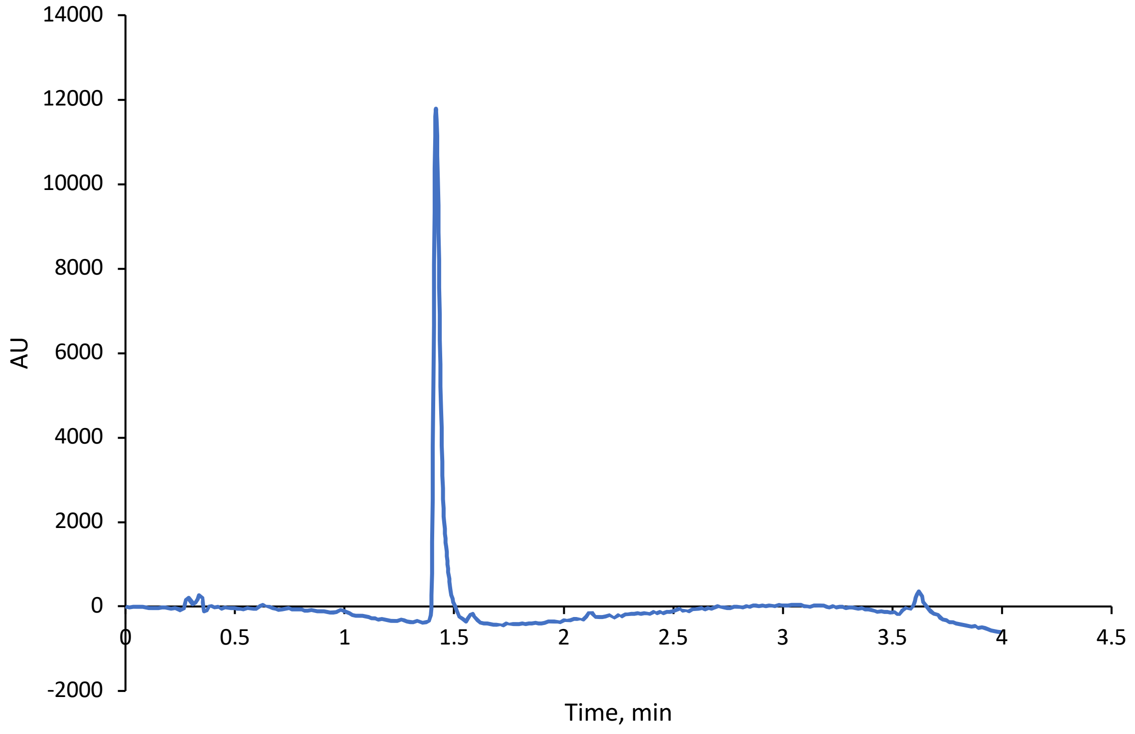

HRMS

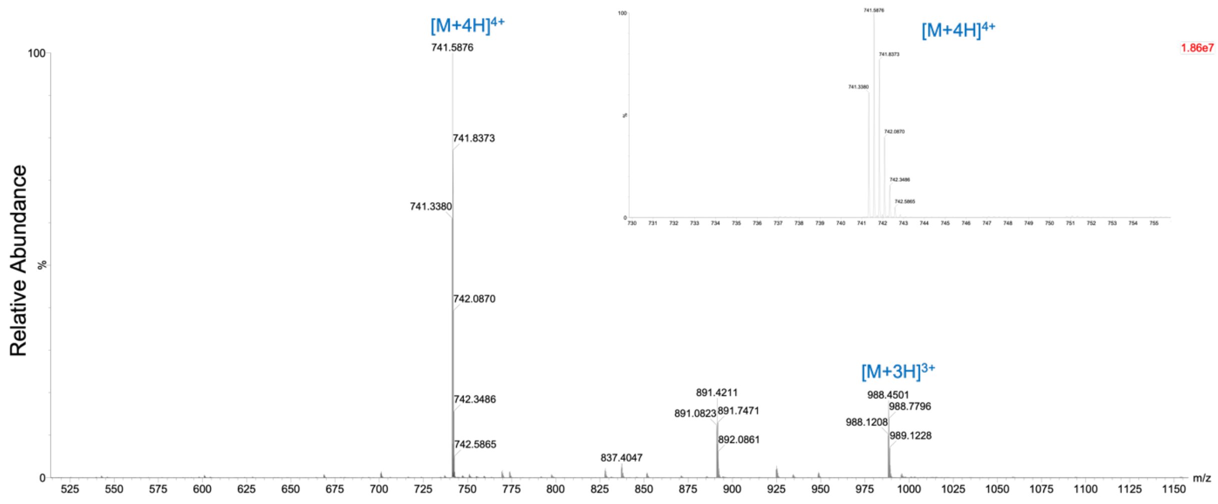
